# Macrophage Xanthine Oxidoreductase Links LPS Induced Lung Inflammatory Injury to NLRP3 Inflammasome Expression and Mitochondrial Respiration

**DOI:** 10.1101/2023.07.21.550055

**Authors:** Mehdi A. Fini, Jenifer A. Monks, Min Li, Evgenia Gerasimovskaya, Petr Paucek, Kepeng Wang, Maria G. Frid, Steven C. Pugliese, Donna Bratton, Yen-Rei Yu, David Irwin, Michael Karin, Richard M. Wright, Kurt R. Stenmark

**Affiliations:** Department of Medicine, Division of Pulmonary and Critical Care; Department of Pediatrics, Division of Pulmonary and Critical Care; Cardiovascular Pulmonary Research Laboratories; Department of Obstetrics and Gynecology; Department of Medicine, Division of Neurosurgery, University of Colorado Denver, Anschutz Medical Campus, Aurora, Colorado, USA; Department of Immunology, UConn Health, Farmington, CT, USA; Department of Medicine, Division of Pulmonary and Critical Care, UPenn, PA, USA; National Jewish Medical Research Center, Denver, Colorado, USA; Department of Pharmacology, University of California San Diego, California, USA

## Abstract

Acute lung injury (ALI) and the acute respiratory distress syndrome (ARDS) remain poorly treated inflammatory lung disorders. Both reactive oxygen species (ROS) and macrophages are involved in the pathogenesis of ALI/ARDS. Xanthine oxidoreductase (XOR) is an ROS generator that plays a central role in the inflammation that contributes to ALI. To elucidate the role of macrophage-specific XOR in endotoxin induced ALI, we developed a conditional myeloid specific XOR knockout in mice. Myeloid specific ablation of XOR in LPS insufflated mice markedly attenuated lung injury demonstrating the essential role of XOR in this response. Macrophages from myeloid specific XOR knockout exhibited loss of inflammatory activation and increased expression of anti-inflammatory genes/proteins. Transcriptional profiling of whole lung tissue of LPS insufflated XOR^fl/fl//LysM-Cre^ mice demonstrated an important role for XOR in expression and activation of the NLRP3 inflammasome and acquisition of a glycolytic phenotype by inflammatory macrophages. These results identify XOR as an unexpected link between macrophage redox status, mitochondrial respiration and inflammatory activation.

## INTRODUCTION

Acute lung injury (ALI) and the more severe acute respiratory distress syndrome (ARDS) remain highly prevalent inflammatory lung disorders in which innate immunity and inflammation are critical mediators of disease [1–4]. ALI can trigger acute and prevalent lung inflammation and fibrosis through activation of the NLRP3 inflammasome and subsequent secretion of IL-1β [5]. Recent studies have shown that targeting NLRP3 may be beneficial in ALI[6]. However, it remains unclear how ALI results in upregulation of NLRP3 and its activation. One possibility is the contribution of XOR . As a source of ROS, RNS, and uric acid (UA) the enzyme Xanthine Oxidoreductase (XOR) has been found to play a critical role in diverse inflammatory disorders[7]. Pharmacological inhibitors of XOR reduced ROS generation and inflammation in both humans and animal models, and they have achieved important clinical utility for the treatment of a wide range of inflammatory disorders [8]. Inhibition of XOR was found to modulate ALI *in vivo* [9–11], to regulate leukocyte adhesion *in vivo* [12–14] and thus reduce neutrophil recruitment to an inflammatory site *in vivo* [9, 15]. Pharmacological inhibition of XOR specifically within macrophages modulated lung inflammation and neutrophil recruitment following adoptive transfer of inflammatory cells [9] and suppressed MCP-1 expression in LPS treated human THP-1 macrophages *in vitro* as well as MCP-1 expression in peritoneal macrophages from LPS injected mice [16].

We thus sought to test the hypothesis that myeloid XOR is a critical mediator of inflammation and innate immunity that could promote ALI/ARDS. While germline knockout of XOR in heterozygous mice (XOR+/-) has led to important insights concerning its role in lactation and mammary gland biology [17], adipocyte differentiation [18], and metabolic regulation [19, 20], homozygous germline knockout mice (XOR-/-) exhibit early neonatal lethality and cannot be used for analysis of inflammatory disorders that are routinely performed on adult mice. For this reason, we sought to develop a conditional cell specific knockout of XOR using a Cre recombinase/LoxP deletion strategy. Here we describe the use of these mice to elucidate the function of myeloid specific XOR on the development of ALI in mice. Analysis of macrophage specific XOR ablation identified an unexpected dual role of XOR in both LPS induced inflammasome expression and modulation of mitochondrial oxidative phosphorylation (OXPHOS).

## RESULTS

### Generation of Conditional XOR Knockout Mice

To produce a conditional myeloid specific knockout of XOR, we generated mice carrying LoxP sites flanking exons E3 and E4 as illustrated (Figures 1A-D). XOR activity and immuno-histochemical staining were unaltered by incorporation of LoxP sites compared to parental C57BL/6 mice (Figures 1E,F), while ectopic expression of Cre recombinase cDNA produced efficient ablation of XOR activity in purified TGE (thioglycolate-ellicited) peritoneal macrophages from XOR^fl/fl^, but not C57BL/6 control mice (Figure 1G). To knockout XOR specifically in myeloid lineage cells, including neutrophils and macrophages, we crossed XOR^fl/fl^ mice with C57BL/6:LysM-Cre mice to create the XOR^fl/fl//LysM-Cre^ strain as described below.

**Figure 1.**
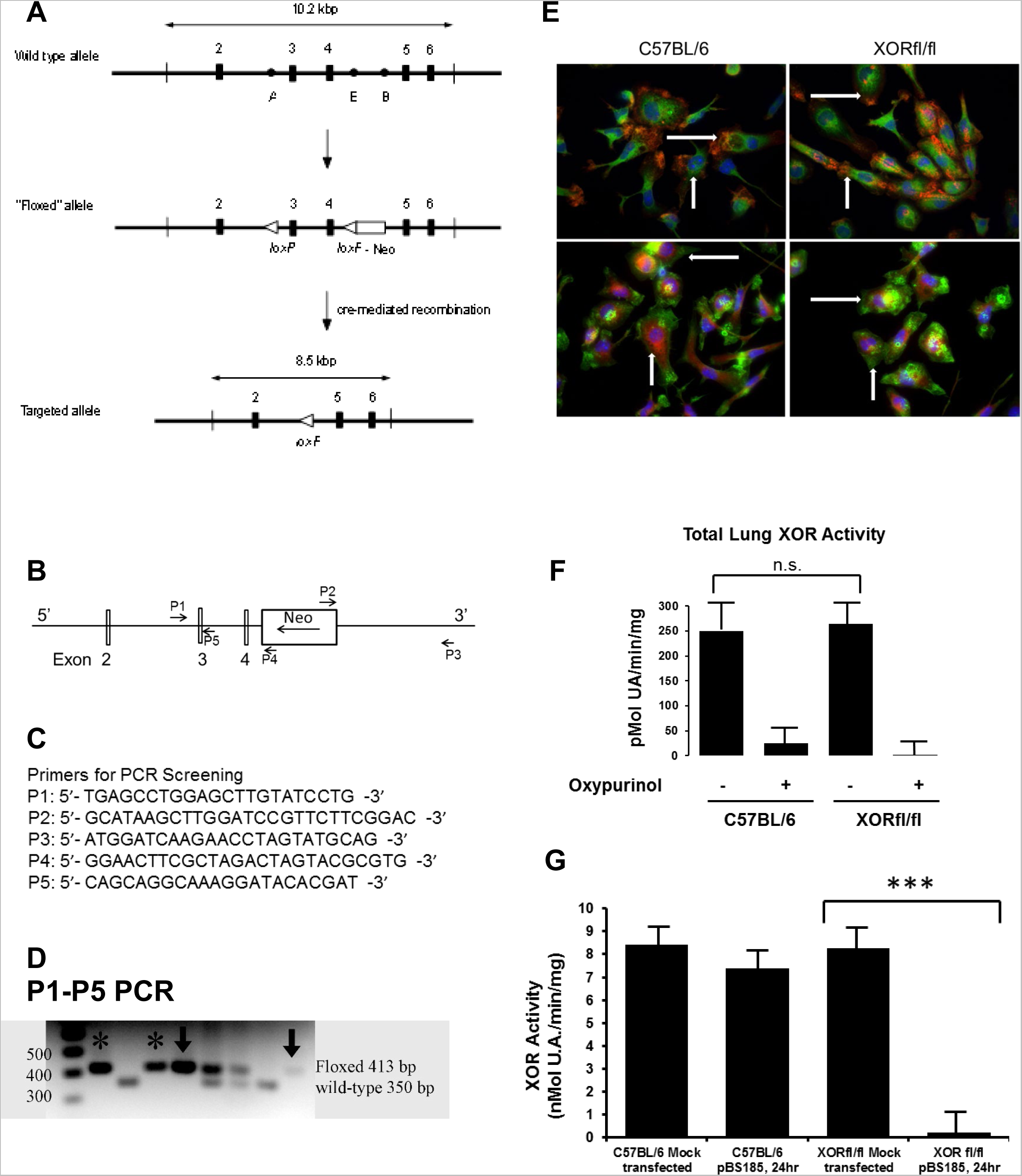
Generation of Conditional XOR Knockout Mice. (A) Map of XOR genomic DNA showing placement of LoxP sites flanking E3 and E4. Cre recombinase mediated recombination will generate a shift in the XOR reading frame resulting in failure to synthesize more than 42 amino acids of 1,338 present in the mature protein. Heterozygous mice carrying the floxed XOR allele were bred to homozygosity, and two separate founder strains of XORfl/fl mice were selected after tail-tip DNA PCR analysis. (B) Genomic PCR screening strategy showing location of PCR primers. (C) PCR primer sequences. (D) P1-P5 PCR gel analysis identifies both the wild type and “floxed” alleles of E4 at 413 and 350 bp respectively. (E) We observed no difference in XOR immunohistochemical staining between C57BL/6 and XORfl/fl TGEM. C57BL/6 and XORfl/fl mice were injected intraperitoneally with 1 ml of 40mg/ml thioglycolate to elicit leukocyte infiltration. After 72 hrs cells were harvested and macrophages purified by adhesion in RPMI 1640 with 10% FBS. Non-adherent cells were rinsed off and the adherent TGEM were stained for XOR (red), nuclei (DAPI in blue), and cell shape actin (green). Two representative high field photos are shown for each strain (top and bottom respectively). Arrows show cytoplasmic stain for XOR. (F) Oxypurinol (150 uM) inhibited total lung XOR activity was determined as described for parental C57BL/6 and XORfl/fl mice using whole lung protein extracts. Data show mean and S.D. of six mice in each group. (G) Ectopic Cre recombinase expression abrogates XOR activity in XORfl/fl but not in C57BL/6 TGEM. Cells were plated as in E and after 24 hours washed cells were transfected with pBS185 or mock transfected using empty vector (pBluescript) at a density of 1.0E6 cells/well. pBS185 is an expression plasmid expressing WT bacteriophage P1 Cre recombinase from the constitutive CMV promoter. XOR activity was determined from six independent transfections 24 hrs after transfection. Data show mean and SE and were analyzed by Students t-test (***, p<0.001).

### Myeloid Specific XOR Knockout Reduced LPS Induced Acute Lung Inflammation and reduced XOR activity and Superoxide Generation *In vivo*

To test the effect of myeloid specific XOR knockout in a common endotoxin model of ALI we insufflated E. coli lipopolysaccharide (LPS) directly into the airway of XOR^fl/fl^ and XOR^fl/fl//LysM-Cre^ mice. After 48 hrs mice were sacrificed and the impact of myeloid specific XOR knockout on inflammation and ALI was determined.

Histological analysis of lung tissue from XOR^fl/fl^ mice demonstrated robust accumulation of inflammatory cells whereas LPS insufflated XOR^fl/fl//LysM-Cre^ littermates exhibited marked reduction in inflammation and inflammatory cell accumulation (Figure 2A). Recovery of macrophages and neutrophils in the bronchoalveolar lavage (BAL) was significantly reduced in XOR^fl/fl//LysM-Cre^ mice (Figure 2B), and both free and adherent macrophage F4/80 staining was markedly reduced in XOR^fl/fl//LysM-Cre^ mice (Figure 2C). Macrophages recovered in the BAL after LPS insufflation were predominantly F4/80+/CD11b+ (15.6% vs 1.12% CD11c+) in XOR^fl/fl^ mice but predominantly F4/80+/CD11c+ (23.7% vs 4.95% CD11b+) in XOR^fl/fl//LysM-Cre^ mice (Figure 2D).

**Figure 2.**
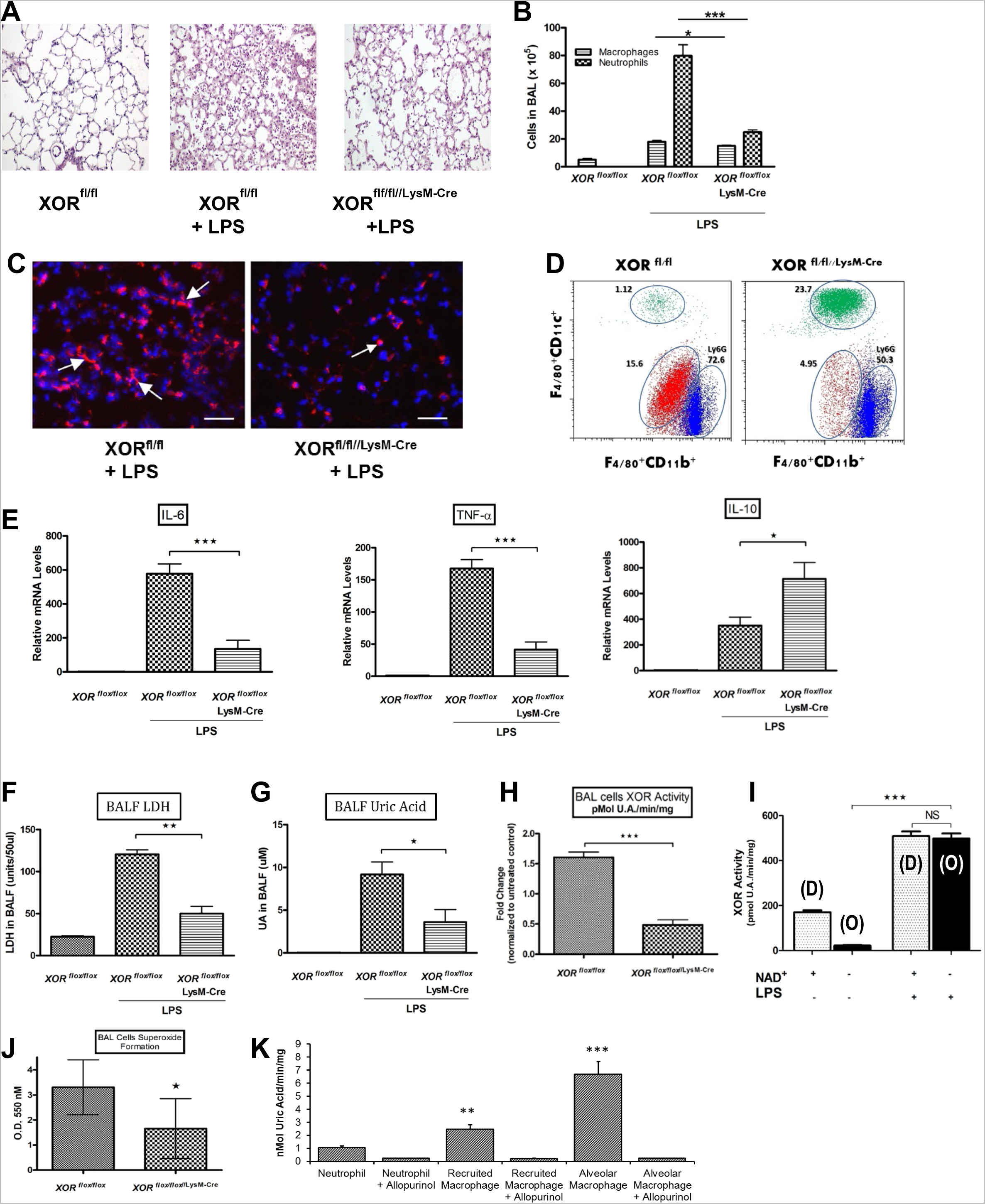
Myeloid Specific XOR Knockout Reduces BAL Cell XOR Activity, Superoxide Generation and Lung Inflammation In LPS Insufflated Mice. (A) 48 hrs after insufflation of either LPS (5.0 mg/kg) or saline (control), mice were sacrificed, lungs perfused blood-free, and prepared for histology. Tissue sections were stained by H&E, photographed, and representative specimens are shown from six independent mice/group. (B) BAL cell analysis 48 hrs after insufflation of either LPS or saline. Macrophage and neutrophil counts were obtained using Wright’s stained BAL cytospins. Data from six mice in each group were analyzed by ANOVA and showed p<0.0001 for both neutrophils and macrophages in XORfl/fl mice comparing with and without LPS insufflation. *, p,0.05 and ***, p<0.001 for macrophage and neutrophil counts with LPS insufflation, respectively. (C) Lung tissue cryosections (5 um) were stained with DAPI (blue) to identify nuclei and the macrophage specific marker F4/80 (red), examined by confocal microscopy, and colors merged from the same fields (scale bar = 50 um). Arrows indicate intra-alveolar free and adherent macrophages. (D) BAL cells from XOR^fl/fl^ and XOR^fl/fl//LysM-Cre^ littermates previously insufflated with LPS were washed and stained for FACS analysis using antibody for F4/80+, CD11b+, and CD11c+. Neutrophils were identified by staining with antibody for Ly6G (FACS, n= 6/6). (E) BAL cells from LPS insufflated XOR^fl/fl^ and XOR^fl/fl//LysM-Cre^ littermates were analyzed by quantitative RT-PCR for inflammatory cytokines IL-6, TNFa and IL-10 (n=6 mice/group). (F) LDH was quantitated in the lung cell-free BAL fluid (BALF) from mice used in panel B (n=6 mice/group). (G) Uric acid was measured in the lung cell-free BALF from mice used in panel B as well (n=6 mice/group). (H) BAL cell XOR activity was determined from XORfl/fl and XORfl/fl//LysM-Cre mice 48 hrs after LPS insufflation. Data show mean and SE (***, p< 0.001) of six independent mice in each group. (I) XOR D-Form and O-Form were determined in BAL cells from saline or LPS insufflated XORfl/fl mice by inclusion or omission of NAD+ from the XOR assay (XOR assay, n= 6/6). (J) BAL cell superoxide generation was determined using cytochrome c reduction for XORfl/fl and XORfl/fl//LysM-Cre mice 48 hrs after LPS insufflation (right panel). Data show mean and SE (*, p< 0.05) of six independent measurements. (K) XOR activity in FACS sorted XORfl/fl BAL cells. Total BAL cells were sorted by FACS following LPS insufflation as described in Figure S2A. Data show the mean and S.E. for three independent cell sortings in which BAL cells from 5 mice per group were pooled. Total NAD+ dependent XOR activity was determined in the presence or absence of Allopurinol. **, p< 0.05, ***, p< 0.001 determined by ANOVA relative to uninhibited neutrophil activity (lane 1).

BAL cells from LPS insufflated XOR^fl/fl^ mice showed marked increase in mRNA for inflammatory cytokines IL-6 and TNFα that was significantly reduced (P<0.001) in XOR^fl/fl//LysM-Cre^ mice while at the same time showing significant increase in anti-inflammatory IL-10 mRNA in the XOR^fl/fl//LysM-Cre^ mice (Figure 2E). While LPS insufflation increased BALF LDH levels by more than 5 fold in XOR^fl/fl^ mice, this was significantly reduced in XOR^fl/fl//LysM-Cre^ mice (Figure 2F). Furthermore, the increased cell free UA recovered in the BALF in mice insufflated with LPS was reduced over 70% in XOR^fl/fl//LysM-Cre^ mice (Figure 2G).

BAL cell XOR activity (Figure 2H) was significantly reduced in XOR^fl/fl//LysM-Cre^ mice compared to XOR^fl/fl^ mice 48 hrs after LPS insufflation. Further we determined that XOR activity in XOR^fl/fl^ BAL cells was largely in Oxidase (O-Form) (>90%) following LPS insufflation but predominantly Dehydrogenase (D-Form) (80%) following saline insufflation (Figure 2I), consistent with XOR as a source of ROS [21, 22]. LPS-induced BAL cell superoxide levels were significantly reduced in XOR^fl/fl//LysM-Cre^ cells versus XOR^fl/fl^ cells (Figure 2J), while lung tissue nitrotyrosine staining was reduced to background levels in XOR^fl/fl//LysM-Cre^ mice compared to XORfl/fl mice (Figure S2A).

To determine which cells in the BAL expressed XOR, cells recovered by lavage from XOR^fl/fl^ mice following LPS insufflation were sorted by FACS into neutrophil (PMN), recruited monocyte/macrophages, and resident alveolar macrophage populations (Figure S1A). Very low levels of Allopurinol inhibited XOR activity were detected in the PMN fraction, while recruited and resident alveolar macrophages together accounted for 94% of the Allopurinol inhibited XOR activity (Figure 2K). Resident macrophages strongly expressed XOR (Figure 2K). Macrophage survival determined by SRB stain was unaltered by XOR genetic ablation (Figure S1B). Pharmacological inhibition of XOR with Allopurinol also had no apparent effect on either macrophage apoptosis (Annexin/PI stain) or survival (Figure S1C), while it blocked M-CSF and GM-CSF induced surface expression of CD11a*^Hi^* and CD11b*^Hi^* in BAL macrophages *in vitro* (Figure S1D,E). Together, these data demonstrate a critical role for myeloid XOR in LPS induced ALI *in vivo*.

### Transcriptional Profiling of Lung tissue in Myeloid Specific XOR Knockout Reveals Broad Effects On LPS Induced ALI *In vivo*

To determine the specific effects of LysM-Cre mediated XOR knockout, Transcriptional profiling of whole lung tissues was performed 8 and 24 hrs after LPS insufflation [23] and microarray data were generated for XOR^fl/fl^ and XOR^fl/fl//LysM-Cre^ mice exposed to LPS and scored for genes up- or down-regulated by myeloid specific XOR knockout.

Genes down-regulated by >1.5 fold 8 and 24 hrs after LPS insufflation by XOR deletion generated a data set of 624 and 564 genes respectively. Genes showing both high enrichment scores (10.6 to 4.31) and low p-value scores (1.0E-12 to 6.3E-5) were sorted by functional category (Tables S1, S3). Inflammatory mediators down-regulated by myeloid specific XOR knockout at 8 hrs included TNFα, IL-1β, iNOS, HIF-1α, α and β alleles of MIP genes 1, 2, and 3, TLR2, and TLR6 (Table S1). Inflammatory mediators down-regulated by myeloid specific XOR knockout at 24 hours demonstrated to overlap with those detected at 8 hours included IL-1β, IL1-R2, CCR1/MIP1αR, CCL4/MIP1β, and TLR6 (Table S3). Evidence for XOR involvement in inflammasome expression was shown by down-regulation of NALP3/NLRP3, IL-1β, and IL-18rap/IL18-Rβ at both 8 and 24 hrs after LPS insufflation. Genes mediating development and function of innate immunity that were down-regulated at 24 hours by myeloid specific XOR knockout included Pou2/Oct2, CD300/CD300a, FOS/AP1, CD11B, S100A9, and PI3K-δ and its associated proteins, consistent with a key role for myeloid XOR in promoting lung inflammation (Table S1 and S3).

Genes up-regulated at 8 and 24 hrs exhibited enrichment scores of 3.08 to 18.05 and p-values from 8.0E-5 to 1.3E-40 (Table S2 and S4). The most significant effect of XOR knockout was observed for genes encoding proteins of OXPHOS including cytochrome b, cytochrome c oxidase (COX) genes 1, 2, 3, and 7A1 as well as NADH dehydrogenase genes ND1, ND2, ND3, ND4, ND5, and ND6. Three genes encoding subunits of the F1/F0 ATPase were also significantly up regulated. 22 mitochondrially encoded t-RNA gene (Trn) and ribosomal RNA genes (Rnr1, Rnr2) mediating mitochondrial translation were also significantly elevated by myeloid specific XOR knockout. Additionally, the mitochondrial transcription factor mtTF2B/TF2BM was also significantly induced at 24 hours by XOR knockout (p = 1.3E-40). No effect on mt*Tfam* was observed. Genes encoding immunoglobulins involved in antigen binding, cell surface mediated signaling, adhesion/migration, and inflammation were significantly up-regulated by myeloid XOR knockout at 24hrs (Table S4). Collectively, these data demonstrated that dual role played by XOR in both suppressing expression of genes encoding mitochondrial OXPHOS while at the same time promoting expression of inflammatory mediators typical of the M(LPS) phenotype [24] including NLRP3 and IL-1β; processes that are lost by myeloid specific XOR knockout.

### Macrophage XOR Contributes To LPS Induced Inflammasome Expression

In agreement with the microarray analysis data (Tables S1-4), we also observed significant decreases in NLRP3, IL- 1β, and TNFα mRNA in XOR^fl/fl//LysM-Cre^ mice compared to XOR^fl/fl^ mice when whole lung RNA from saline or LPS insufflated mice was screened by quantitative RT-PCR analysis (Figure S2B-D). Macrophages purified from the BAL of LPS insufflated XOR^fl/fl^ and XOR^fl/fl//LysM-Cre^ mice also revealed significantly decreased mRNA expression for NLRP3 and IL-1β (Figure S2E,F). The decrease in mRNA for IL-18 followed a similar trend but did not reach statistical significance (Figure S2G). Both NLRP3 and IL-1β protein levels were also reduced in the purified macrophages from XOR^fl/fl//LysM-Cre^ mice compared to XOR^fl/fl^ mice (Figure 3A). Inflammatory cytokine expression in the cell free BALF was analyzed by ELISA and western blot which demonstrated significantly decreased IL-1β and IL-18 secretion consistent with the reduced expression of IL-1β mRNA in the BAL macrophages of LPS insufflated XOR^fl/fl//LysM-Cre^ mice (Figure 3B). We also observed marked reduction in pro-caspase-1 and mature p10/p20 caspase-1 proteins in both XOR^fl/fl//LysM-Cre^ lung tissues (Figure S2H) and in whole cell lysates of macrophages purified from the BAL of LPS insufflated mice (Figure 3C).

**Figure 3.**
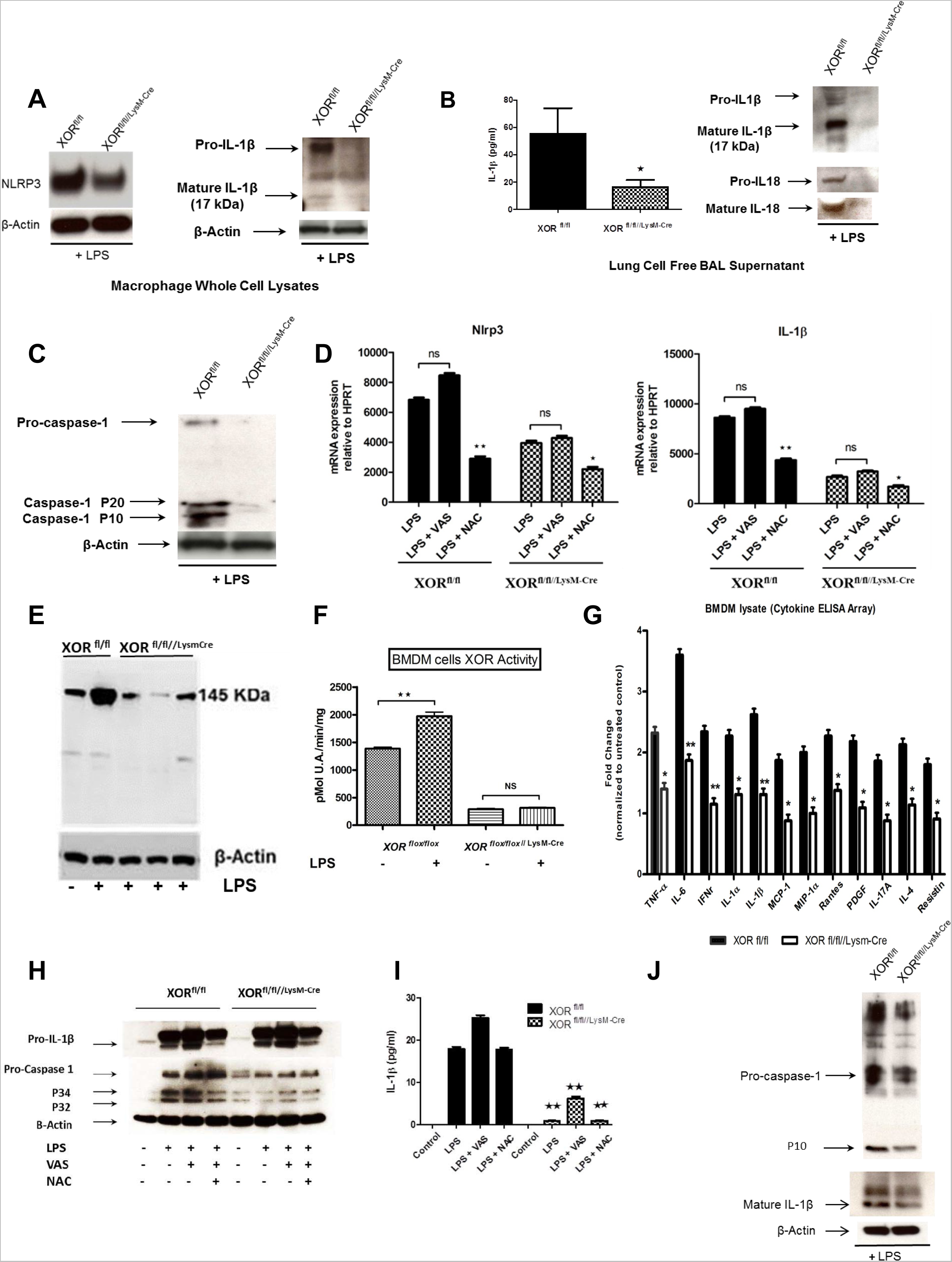
Macrophage XOR Contributes To LPS Induced Inflammasome Expression and Inflammarotory Activation. (A) Whole cell lysates of purified macrophages from saline or LPS insufflated XORfl/fl and XORfl/fl//LysM-Cre mice were analyzed by western blot for protein levels of Nlrp3 and IL-1β (A, n=6/6). (B) Cell free BAL fluid collected 48 hrs after saline or LPS insufflation in XORfl/fl and XORfl/fl//LysM-Cre mice was analyzed by ELISA for expression of secreted IL-1β (left panel) and by western blot for protein levels of both IL-1β and IL-18 (right panel). Data were analyzed as in (B); **, p<0.02. (IL-1b, n=8/8) (C) Whole cell lysates of BAL macrophages purified from XORfl/fl and XORfl/fl//LysM-Cre mice insufflated with LPS 48 hrs before were analyzed by western blot for expression of caspase-1 p10 using HRP conjugated antisera against caspase-1(n=6/6). (D) qRT-PCR of Nlrp3 and IL-1β mRNA from macrophages purified from XORfl/fl and XORfl/fl//LysM-Cre mice 48 hrs after insufflation of LPS or saline. Purified macrophages were cultured in vitro in the presence of 1.0 mM NAC or 10.0 uM VAS2870 (Sigma, SML0273). After 24 hrs cells were harvested and analyzed by qRT-PCR for expression of Nlrp3 and IL-1β. Data show the mean and SE of six independent measurements with **, p< 0.02 relative to LPS alone; *, p< 0.05 relative to LPS alone. (E) Western immunoblot analysis of BMDM XOR protein from XORfl/fl and XORfl/fl//LysM-Cre mice treated with 100 nM LPS/ 24hr, three independent preparations of BMDM. (F) BMDM from XORfl/fl and XORfl/fl//LysM-Cre mice were treated with LPS (100 ng/ml) or saline (control) and after 24 hrs cells were lysed and XOR activity determined from six independent wells as in panel B. (G) Mouse cytokine ELISA array (Signosis, Inc., Santa Clara, CA, USA) was used on whole cell lysates from these cells as described by the supplier. Data were normalized to the untreated WT control and shown as the mean and S.E. XOR^fl/fl^ and XOR^fl/fl//LysM-Cre^ data were analyzed by Students t-test (*, p<0.05; **, p<0.02) (ELISA, n=8/8). (H) Western immunoblot analysis of IL-1β and Caspase-1 in whole cell lysates used in panel G. (I) IL-1β ELISA of the cell-free culture medium from cells used in panel I. Data show the mean and SE of four replicas (**, p< 0.02) when data were analyzed by pair-wise Student’s t-test comparing LPS vs LPS, LPS/VAS vs LPS/VAS, and LPS/NAC vs LPS/NAC. (J) Whole cell lysates from purified BAL macrophages generated as in panel D were analyzed for expression of IL-1β and caspase-1 by western blot.

To determine if XOR derived ROS mediated expression of inflammasome components, purified BAL macrophages were cultured in the presence of the ROS scavenger N-acetylcysteine (NAC) or VAS2870, a selective inhibitor of NADPH oxidase activity (NOX) and ROS generation. After 24 hrs cells were harvested and analyzed by qRT-PCR for expression of Nlrp3 and IL-1β. While VAS2870 had no significant effect on Nlrp3 or IL-1β expression, both NAC and XOR knockout significantly reduced expression of both Nlrp3 and IL-1β (Figure 3D). The combination of NAC and the XOR knockout revealed a small but significant contribution to LPS induced Nlrp3 and IL-1β that was not derived from XOR generated ROS alone. Nonetheless, these data confirm the principal role of XOR derived ROS in the expression of LPS induced Nlrp3 and IL-1β, but do not support a primary role for the NADPH oxidase (Figure 3D).

To determine the effects of XOR ablation on macrophage cells other than those resident or recruited to the lung alveolar space in the IT LPS model, XOR^fl/fl^ and XOR^fl/fl//LysM-Cre^ mice were used to evaluate the contribution of XOR to activation of thioglycollate-elicited peritoneal macrophages (TGEM). BMDM exhibited >90% positive staining for the macrophage marker F4/80 (Figure S3A), and >95% viability following M-CSF differentiation (Figure S3B). XOR activity and protein level were reduced by >70% in LPS treated BMDM and TGEM from XOR^fl/fl//LysM-Cre^ mice compared to XOR^fl/fl^ littermates (Figure 3E,F; Figure S4A,B). While F4/80 immunoreactivity (Figure S3A) and cell survival were unaffected by XOR knockout in XOR^fl/fl//LysM-Cre^ cells (FigureS3B).

Treatment of BMDM with 100 ng/ml E. coli LPS for 24 hrs increased XOR immunoreactivity that was reduced to near background in XOR^fl/fl//LysM-Cre^ compared to XOR^fl/fl^ BMDM (Figure 3E). XOR activity, and both basal and LPS induced XOR activity was blocked by XOR knockout in XOR^fl/fl//LysM-Cre^ mice (Figure 3F). LPS treatment did not reduce survival of M-CSF differentiated BMDM in either XOR^fl/fl^ or XOR^fl/fl//LysM-Cre^ cells (Figure S3B). The inflammatory marker Galectin-3 showed marked stimulation by LPS in XOR^fl/fl^ derived BMDM that was reduced to basal levels in XOR^fl/fl//LysM-Cre^ cells (Figure S3D). We observed between 2 and 4 fold increase in several inflammatory cytokines in BMDM from XOR^fl/fl^ mice that was significantly reduced (TNFα, IL-6, IL-1α, IL-1β, RANTES) or blocked completely in XOR^fl/fl//LysM-Cre^ cells after LPS treatment (MCP-1, MIP-1α, IL-17A, Resistin) (Figure 3G).

In contrast to BAL macrophages, western immunoblot analysis of BMDM revealed little or no effect of XOR ablation on intracellular levels of LPS stimulated pro-IL-1β (Figure 3H). We observed a small decrease in mature IL-1β by western blot and ELISA (Figure 3G, J) as well as a 2-fold decrease in both pro- and mature caspase-1 in LPS stimulated BMDM from XOR^fl/fl//LysM-Cre^ mice compared to XOR^fl/fl^ mice (Figure 3H, J). Expression of LPS stimulated intracellular pro-IL-1β protein was insensitive to both XOR ablation and inhibition of NADPH oxidase but was reduced by NAC, suggesting an unknown source of ROS that affected LPS induced BMDM pro-IL-1β (Figure 3H). In contrast, intracellular pro-caspase-1 as well as caspase p10, p34, and p32 were reduced 2 fold by XOR ablation or NAC, but not VAS2870 (Figure 3H). NAC treatment in conjunction with XOR ablation did not further reduce caspase proteins suggesting that XOR derived ROS alone contributed to caspase protein expression. Secreted mature IL-1β was markedly inhibited by macrophage XOR knockout (Figure 3 H,I). Furthermore, release of mature IL-1β was XOR dependent but ROS independent, suggesting a potential role for intracellular UA or metabolic state in IL-1β secretion.

### Macrophage Inflammasome Expression Is Signaled By XOR In Part Through NF-κB

XOR derived ROS could mediate inflammatory activation in part through NF-κB, a central regulator of inflammatory cytokine expression in macrophages [25]. We observed marked reduction in Ser32-phosphorylation of the NF-κB activator IκBα in macrophages from LPS insufflated XOR^fl/fl//LysM-Cre^ mice compared to XOR^fl/fl^ mice (Figure 4A). IκBα phospho-Ser32 activation was further reduced by the NF-κB inhibitor BAY-11-7085 (Figure 4B). We observed that the BAL macrophage NLRP3, pro-IL-1β, and pro-caspase-1 all showed increased protein expression by NFκB inhibition, an effect that was entirely abrogated by XOR knockout (Figure 4C). Decrease in NF-κB p65 and p50 was observed in macrophage whole cell lysates from LPS insufflated mice by XOR ablation compared to XOR^fl/fl^ control mice (Figure 4D).

**Figure 4.**
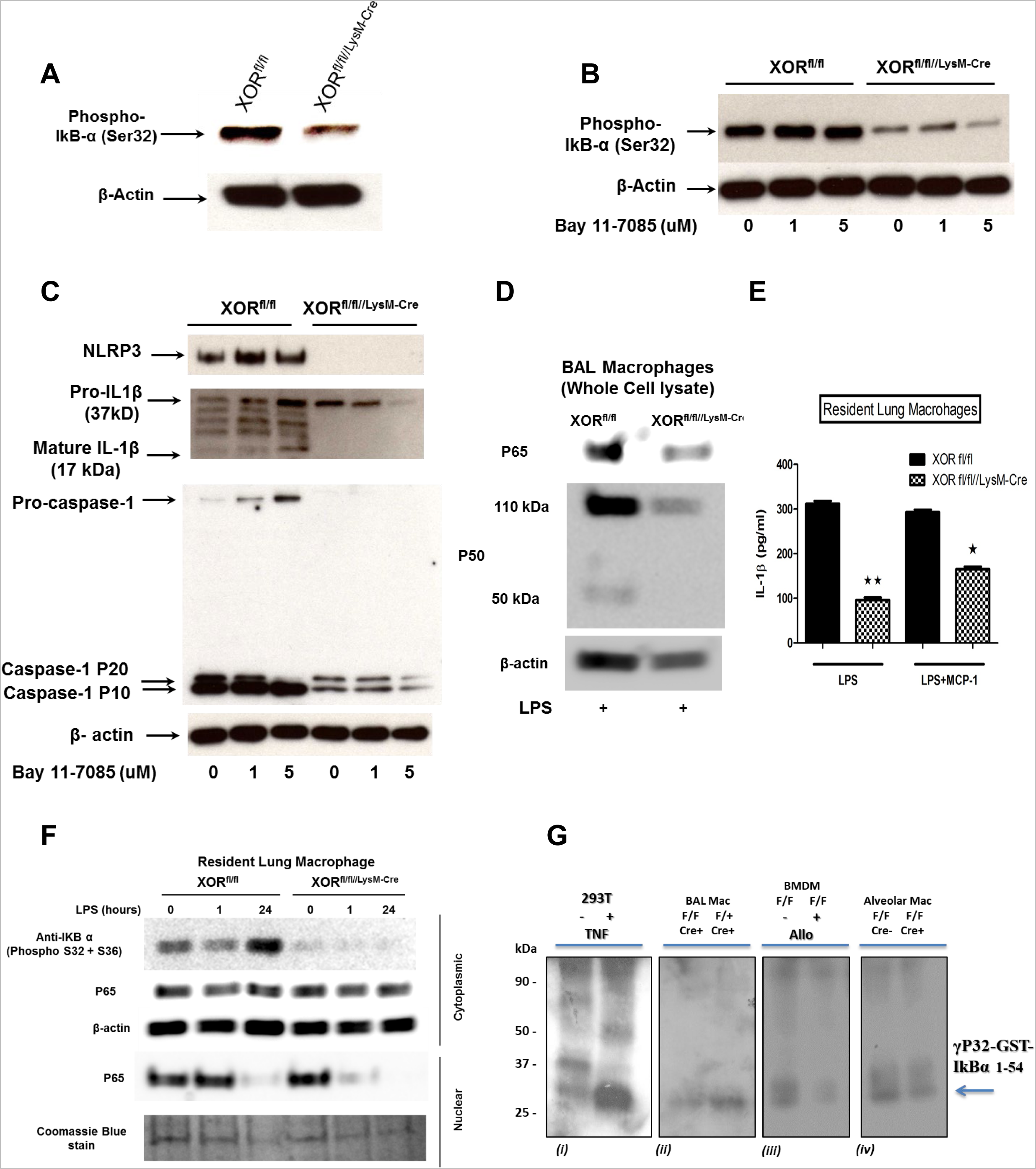
Macrophage Inflammasome Expression Is Signaled By XOR In Part Through NF-κB. (A) Western blot of IκBα phospho-Ser32 using whole cell lysates of BAL macrophages purified from lungs of XORfl/fl and XORfl/fl//LysM-Cre mice insufflated with LPS 48 hrs before. Blot is representative of three independent insufflation experiments and used macrophages pooled from four mice in each group. (B) BAL macrophages from panel A were incubated in vitro with the NF-κB inhibitor BAY-11-7085 at the indicated doses and after one hour lysates were obtained and analyzed by western immunoblot for IκBα phospho-Ser32. (C) Inflammasome components NLRP3, IL-1β, and caspase-1 were analyzed by western blot using the same cell lysates used in panel A. (D) Whole cell lysates of BAL macrophages used in panel A were probed with antisera against NFκB p65 and p50. (E) Resident alveolar macrophages were purified from lungs of untreated mice and exposed to LPS (100 ng/ml) or MCP-1 (20 ng/ml) for 24 hrs. Secreted IL-1β was measured in the cell free medium by ELISA from four independent replicas. **, p<0.02; *, p<0.05. (F) Purified resident alveolar macrophages were cultured in vitro in the presence of (100 ng/ml) for 0, 1, and 24 hrs. Nuclear and cytoplasmic fractions were obtained and analyzed by western immunoblot for IκBα Ser-32/36 phosphorylation p65. Coomassie Blue stain was used to control for protein loading in the nuclear fractions. (G) IKK activity was measured by immunocomplex kinase assay using GST-IκBα(1–54) as a substrate in: (i) lysates of HEK-293 cells treated with 20 ng/ml TNF for 10 min (positive control), (ii) BAL macrophages from heterozygous and homozygous XOR fl/fl//LysmCre+ mice insufflated with LPS after 24 hrs, (iii) BMDMs from LPS- insufflated XORfl/fl mice with and without pretreatment by XOR inhibitor Allopurinol and (iv) alveolar macrophages isolated from C57BL6 and XOR fl/fl//LysmCre+ mice and treated with LPS for 30 min in vitro . Blot is representative of three independent experiments and used macrophages pooled from three mice in each group.

Resident alveolar macrophages from untreated XOR^fl/fl^ and XOR^fl/fl//LysM-Cre^ mice were exposed to LPS or LPS/MCP-1 for 24 hrs *in vitro*. Secreted IL-1β measured in the cell-free culture medium was markedly reduced by XOR ablation (Figure 4E). Consistent with the Ccr2- (negative) status of these cells, no effect was observed by inclusion of the XOR inducer MCP-1. Cytoplasmic and nuclear protein fractions were obtained from alveolar macrophages treated with LPS for 0, 1, and 24 Hrs *in vitro*. We observed that LPS markedly increased cytoplasmic IκBα-Ser32 phosphorylation that was entirely abrogated by XOR ablation (Figure 4F) while nuclear accumulation of NFκB p65 one hour after LPS exposure was also abrogated by XOR ablation (Figure 4F). Furthermore, exposure of human U937 cells to LPS for 4 hrs *in vitro* stimulated nuclear accumulation of both NFκB p65 and p50 that was attenuated by XOR pharmacological inhibition (Figure S5A).

The effect of XOR on IkBa kinase activity was measured using both pharmacological (Allopurinol) and genetic (XOR^fl/fl//LysmCre+^ and XOR^fl/fl^ mice) approaches. In all these in vivo and in vitro approaches, the inhibition of XOR activity consistently resulted in attenuation of IkBa kinase activity (Figure 4G *i-iv*).

### XOR Contributes to the LPS Induced Glycolytic Shift in macrophages

To examine the role of XOR in mitochondrial OXPHOS, we measured levels of lactate and pyruvate in purified BMDM derived from XOR^fl/fl^ and XOR^fl/fl//LysM-Cre^ mice 24 hrs after LPS treatment *in vitro*. A significant reduction in lactate generation as well as free NADH by XOR^fl/fl//LysM-Cre^ macrophages was observed compared to XOR^fl/fl^ macrophages (Figure 5A). The effect of XOR knockout on expression of mitochondrial OXPHOS genes was verified by qRT-PCR of whole lung RNA obtained from saline or LPS insufflated mice. LPS markedly suppressed expression of complex I genes (ND1, ND2, ND3, ND4) and Cytochrome b in XOR^fl/fl^ mice, and this effect was significantly reversed in XOR^fl/fl//LysM-Cre^ mice (Figure 5B, left panel). Purified macrophages isolated from the BAL of mice insufflated with LPS likewise revealed increased expression of these genes in XOR^fl/fl//LysM-Cre^ mice compared to LPS insufflated XOR^fl/fl^ control mice (Figure 5B, right panel). Expression of the mitochondrial transcription factor *Tfam* was not affected by myeloid specific XOR knockout whereas *Tfb2m* exhibited significant two-fold increase in both mRNA and protein level in parallel with expression of both *Pgc-1α* and the mitochondrially encoded OXPHOS gene *CoxII* (Figure 5C,D). We observed that *Tfb2m* expression was increased when XOR^fl/fl^ purified BAL macrophages were cultured in the presence of LPS and NAC. However in XOR knockout macrophages, the LPS effect on *Tfb2m* was not further enhanced by NAC, demonstrating the contribution of WT XOR derived ROS to *Tfb2m* repression (Figure 5E). Together, these data demonstrate the contribution of XOR to LPS induced repression of mitochondrial genes that was in part reversed by myeloid specific XOR knockout.

**Figure 5.**
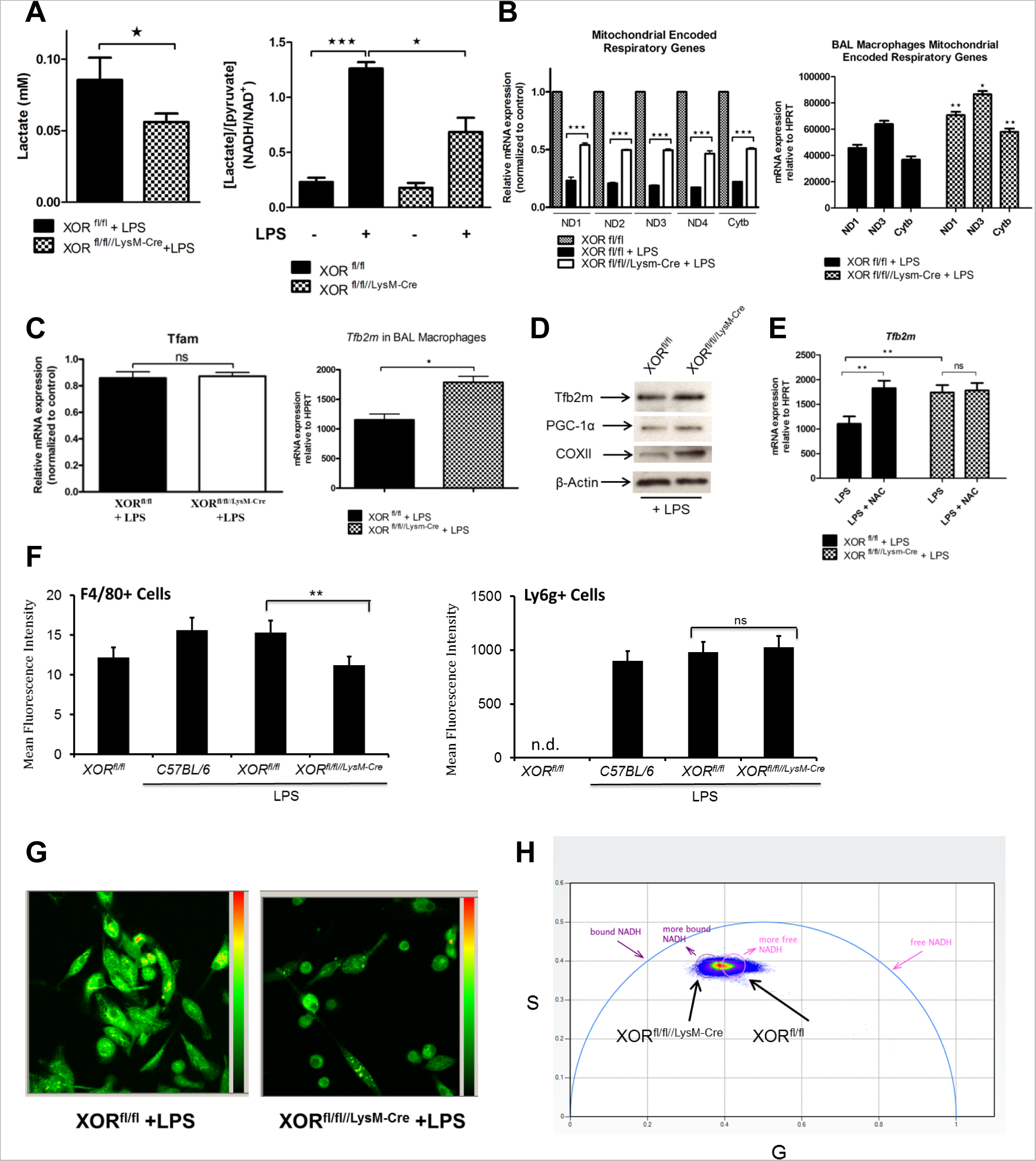
Macrophage XOR contributes to LPS induced glycolytic shift. (A) Levels of lactate and pyruvate were measured in whole cell lysates from BMDM derived from XORfl/fl and XORfl/fl//LysM-Cre mice 24 hrs after LPS (100 ng/ml) treatment in vitro. Data show mean and S.E. of six independent measurements for each group, and data were analyzed by Student’s t-test. (*, p<0.05; ***, p<0.001). (B) Total RNA was obtained from lung tissue 48 hours after LPS insufflation and subjected to qRT-PCR for mitochondrial encoded genes (left panel). RNA was also obtained from macrophages purified from the BAL by adhesion 48 hrs after LPS insufflation and analyzed by qRT-PCR for mitochondrial genes (right panel). Data were analyzed as in panel A with * p<0.05, ** p<0.02, (n=6/6). (C) Total RNA used in panel B was assayed by qRT-PCR for expression of mitochondrial transcription factors Tfam and Tfb2m. Data were analyzed as in panel B. (D) Tfb2m, PGC-1α, and COXII western blot using purified macrophage whole cell extracts.(E) BAL macrophages were purified from LPS insufflated mice, cultured in vitro in the presence of NAC, and after four hours RNA was obtained and analyzed by qRT-PCR for expression of HPRT and Tfb2m. Data show the mean and SE of four independent measurements. (**, p<0.02 by Student’s t-test). (F) Total BAL cells were harvested from LPS insufflated and control mice and stained with MitoSox in the presence of either Alexaflour conjugated anti-F4/80 or Ly6G. Cells were sorted by FACS and mean fluorescence intensity (MFI) from MitoSox determined (**, p < 0.02), (n=6/6). (G) Intrinsic NADH fluorescence of purified BAL macrophages from XORfl/fl and XORfl/fl//LysM-Cre mice insufflated with LPS (5 mg/kg) 48 hrs earlier was detected using Fluorescence Lifetime Imaging (FLIM). (H) Phasor plot analysis of NADH fluorescence shown in G. The Phasor plot shown depicts the presence of the two different NADH species (free and mitochondrial bound) in each individual pixel of both images shown in G. The phasor data contained within the circle show masked pixels of the shorter lifetime values that correspond to free-NADH (pink), whereas the magenta masked pixels show the longer lifetime values correspond to bound-NADH.

To determine if myeloid specific XOR knockout modulated mitochondrial ROS generation, total BAL cells were harvested from LPS insufflated mice and stained with MitoSox in the presence of either Alexaflour conjugated anti- F4/80 (macrophage) or Ly6G (neutrophil) and sorted by FACS. We observed significant (p < 0.02) reduction in mean fluorescence intensity (MFI) from LPS insufflated XOR^fl/fl//LysM-Cre^ macrophages compared to XOR^fl/fl^ or C57BL/6 macrophages that was not observed in the neutrophil population (Figure 5F). Consistent with the decrease in mitochondrial ROS observed in LPS insufflated XOR^fl/fl//LysM-Cre^ mice compared to XOR^fl/fl^, we observed marked decrease in the ratio of free/bound NADH in macrophages from LPS insufflated XOR^fl/fl//LysM-Cre^ mice using Phasor/FLIM analysis compared to macrophages from LPS insufflated XOR^fl/fl^ mice (Figure 5G,H). These data are consistent with the involvement of XOR in promoting a macrophage glycolytic phenotype that shifts toward OXPHOS in XOR^fl/fl//LysM-Cre^ mice and identify XOR as a crucial mediator of the macrophage metabolic state.

### Macrophage XOR contributes to Mitochondrial Respiration and LPS/CoCl2 induced mitochondrial dysfunction

Mitochondrial matrix oxidant burden was assessed via FACS analysis of BMDMs stained with MitoSOX Red. Pretreatment of BMDMs with the pharmacologic XOR inhibitor, Y700, attenuated the LPS-induced MitoSOX Red (Figures 6A). Next, the direct effect of XOR (separate from its effect on the inflammatory microenvironment) on mitochondrial status and function was assessed in macrophage RAW246.7 cells exposed to cobalt chloride (CoCl2) that causes rapid hypoxia in cells with/without Y700 (XOR inhibitor with no ROS scavenging). CoCl2 augmented XOR activity (both D & O Form) in a short time data point (Figures 6B), and increased Mitosox staining, while it was attenuated by Y700 (Figures 6C, S6B). Furthermore, CoCL2 decreased mitochondrial membrane potential as identified by tetramethylrhodamine ethyl ester (TMRE) staining, and this effect was partially reversed by the XOR inhibitor Allopurinol (Figures 6D-E, S6C).

**Figure 6.**
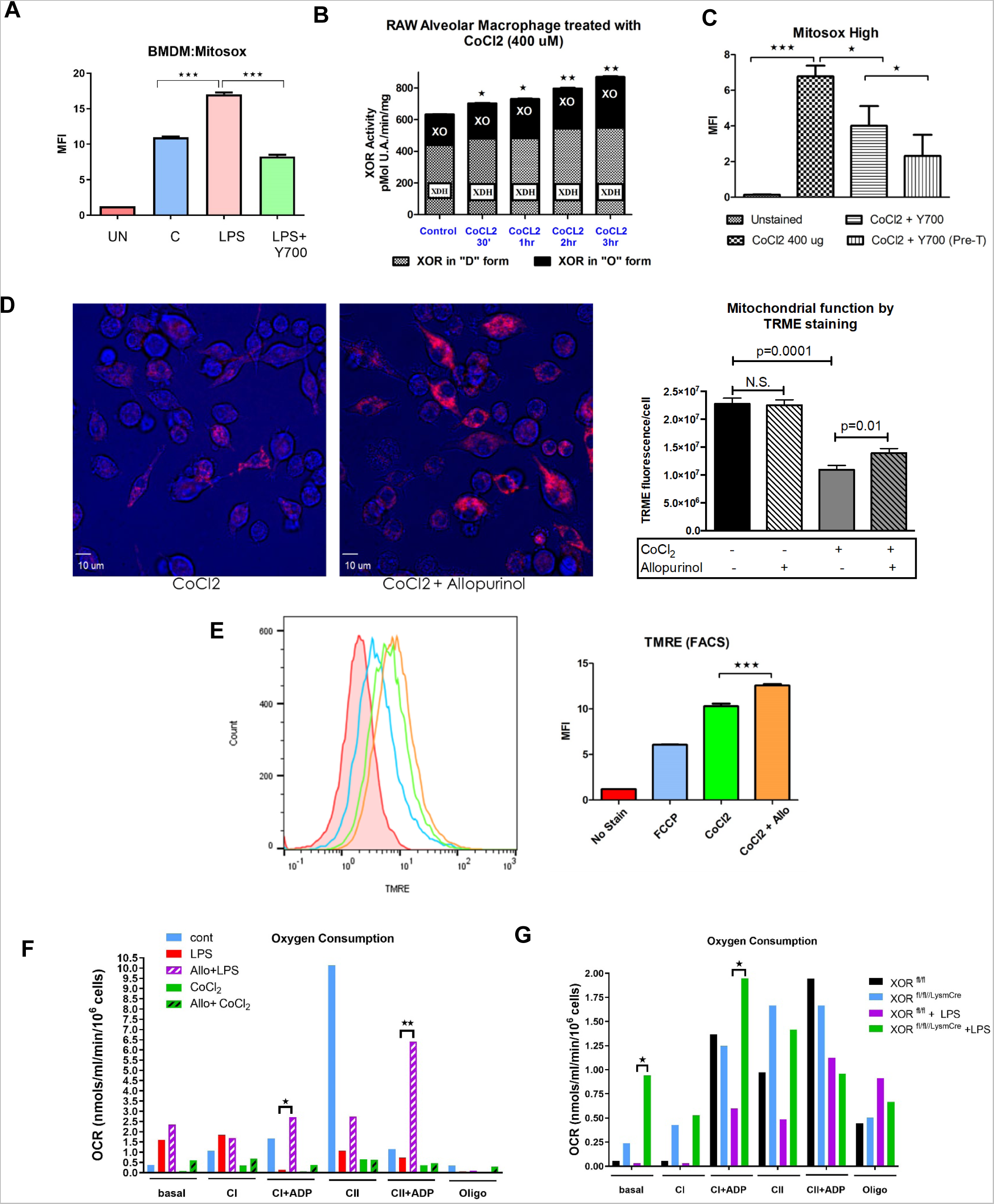
Macrophage XOR Modulates Mitochondrial Respiration in both LPS and CoCl2 induced mitochondrial dysfunction. (A) FACS analysis of BMDM cells stained with MitoSOX Red treated with LPS with and without XOR inhibitor Y700. Data show mean and S.E. of six independent measurements for each group, and data were analyzed by Student’s t- test. (*, p<0.05; ***, p<0.001). (B) XOR activity assay in RAW246.7 cells treated with CoCL2 at different time points(30’-3hrs). Data show the mean and SE of four independent measurements. (*, p< 0.05, **, p< 0.02 determined by ANOVA relative to untreated control (lane 1). (C) FACS analysis of RAW246.7 cells stained with MitoSOX Red treated with CoCL2 with and without XOR inhibitor Y700. Data were analyzed as in panel A with * p<0.05, ** p<0.02, (n=6/6). (D) Analysis of mitochondrial membrane potential using fluorescent microscopy in RAW246.7 cells stained with TMRE and treated with CoCL2 with and without Allopurinol, (Scale bar= 10 um). (E) Analysis of mitochondrial membrane potential with flow cytometer using 488nm laser for excitation and at emission 575 nm, in RAW246.7 cells stained with TMRE and treated with CoCL2 with and without Allopurinol, (n=6/6). (F) Analysis of baseline and maximal mitochondrial respiration rate in BMDM cells treated with CoCL2 (3hrs) and LPS (24hrs) with and without allopurinol, (n=6/6). (G) Analysis of baseline and maximal mitochondrial respiration rate in purified BAL macrophages from XORfl/fl and XORfl/fl//LysM-Cre mice insufflated with LPS (5 mg/kg) 24 hrs earlier using Clark electrode, (n=4/4).

Both CoCL2 and LPS decreased mitochondrial basal and maximal respiration, measured at mitochondrial complex I and II (CI & CII), which was significantly improved by Allopurinol (Figure 6F). Importantly, BAL macrophages isolated from LPS-insufflated XOR LysmCre+ mice, showed differential and improved mitochondrial respiration at basal and CI levels (Figure 6G).

### XOR Myeloid Knockout Inhibits Macrophage Antioxidant Response Through NRF-2β

XOR^fl/fl//LysM-Cre^ macrophages also exhibited significant reduction in both total SOD and catalase levels (Figure 7A, B, C), suggesting that XOR derived ROS contributed to the macrophage antioxidant response. We examined expression of the PRC and NRF family of genes that are involved in control of a wide range of inflammatory genes [26, 27]. In lung tissue extracts from LPS insufflated mice, levels of PRC, NRF-1, and NRF-2β proteins were reduced in XOR^fl/fl//LysM-Cre^ mice compared to XOR^fl/fl^ littermates (Figure 7D, E). The constitutive NRF-2α protein was not significantly affected by myeloid specific XOR knockout (Figure 7D). BAL macrophages purified from lungs of LPS insufflated mice also showed marked loss of NRF-2β by myeloid specific XOR ablation (Figure 7F) as did purified BMDM treated with LPS *in vitro* (Figure 7G). Resident alveolar macrophages purified from untreated XOR^fl/fl^ and XOR^fl/fl//LysM-Cre^ mice were exposed to LPS 0,1, or 24 hrs *in vitro*, and whole cell lysates were analyzed for levels of Keap-1 (Figure 7H). These data revealed marked accumulation of Keap-1 by XOR ablation in LPS treated alveolar macrophages consistent with the reduction in Nrf-2β. These data suggest a mechanism by which XOR derived ROS contribute to Nrf-2β stability through the ROS sensitive Keap-1 regulator protein and suggest that WT XOR derived ROS both activate NFκB dependent inflammatory gene expression while also promoting NRF-2β stability and antioxidant response (Figure 7I).

**Figure 7.**
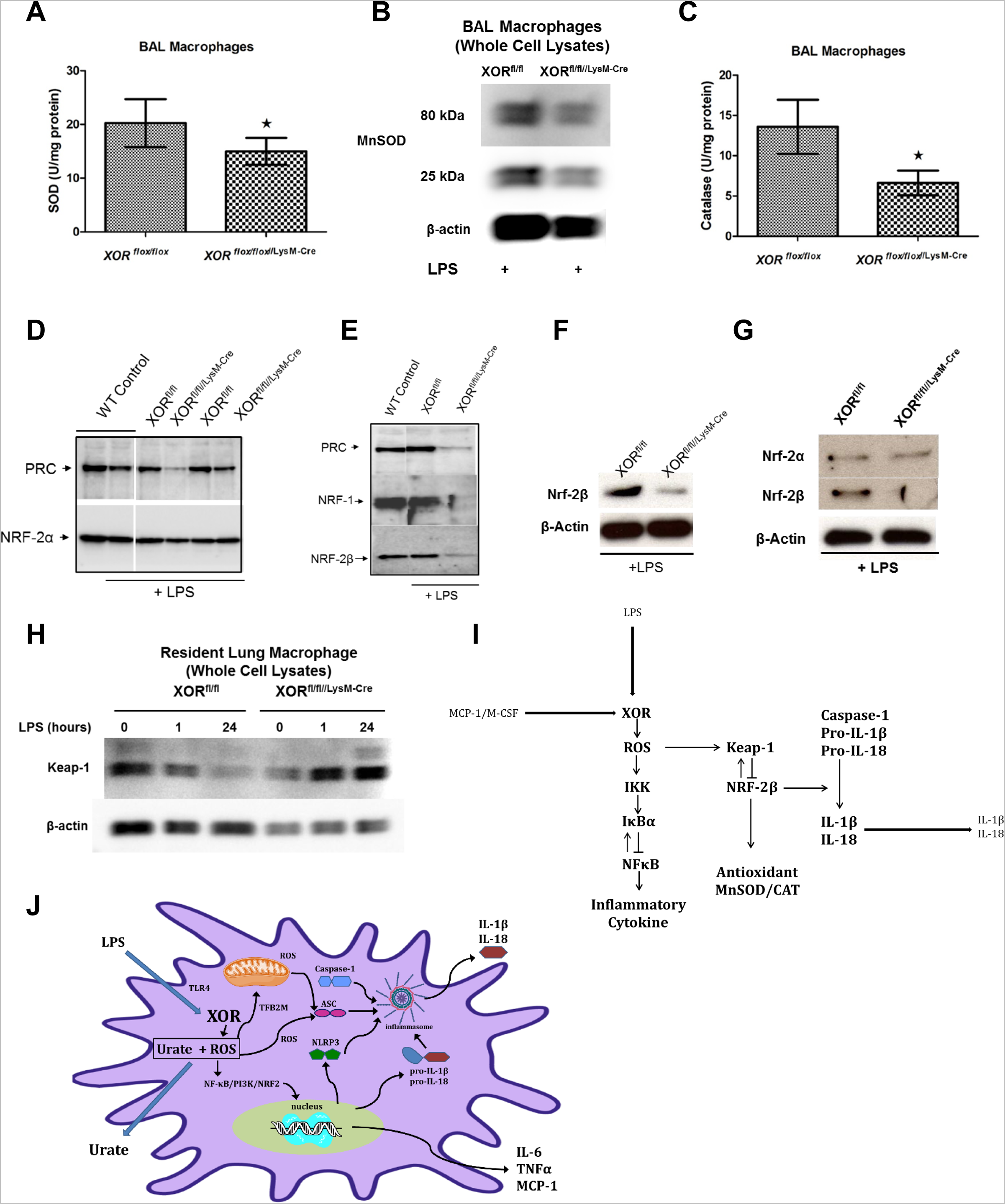
XOR Myeloid Knockout Inhibits Macrophage Antioxidant Response Through NRF-2β. (A) Macrophages purified from the BAL of XORfl/fl and XORfl/fl//LysM-Cre insufflated with LPS 48 before were analyzed for total SOD activity exactly as described using four mice in each group. (*, p<0.05 by Student’s t-test). (B) Western immunoblot of MnSOD using macrophage whole cell lysates from panel A. (C) Macrophages used in panel A were analyzed for total catalase activity. (*, p<0.05 by Student’s t-test, n=4). (D) 48 hrs after insufflation of either LPS (50 mg/kg) or saline (WT control), mice were sacrificed, lungs perfused blood-free, and whole tissue lysates were generated. Western immunoblots were run using antibody against PRC- 1α and NRF-2α. (E) Replica lung tissue extracts were probed with antibody against PRC-1α, NRF-1, and NRF-2β. (F) Western blot of NRF2-β from purified BAL macrophage used in panel A. (G) NRF-2α and NRF2-β western blot of BMDM proteins from cells treated with LPS (100 ng/ml) for 24 hrs in vitro. (H) Purified resident alveolar macrophages from untreated XORfl/fl and XORfl/fl//LysM-Cre mice were cultured in vitro in the presence of LPS (100 ng/ml) for 0, 1, and 24 hrs. Whole cell lysates were obtained and analyzed by western immunoblot for Keap-1, (n=4/4). (I) Illustration showing the proposed mechanism by which macrophage XOR derived ROS contribute to both inflammatory cytokine expression and inflammasome mediated IL-1β and IL-18 maturation. Present data support a mechanism by which XOR derived ROS both promote NF-κB mediated inflammatory cytokine expression while also contributing to Nrf-2β stability and maturation of IL-1β and IL-18. ROS modified Keap-1 is postulated to disrupt Keap-1/Nrf-2β association, inhibit Nrf-2β ubiquitination and degradation, permitting Nrf-2β stabilization and nuclear activation. XOR ablation would both inhibit NF-κB activation and inflammatory cytokine expression as well as reduce Nrf-2β stability leading to reduced maturation of IL-1β and IL-18.

## DISCUSSION

This study identified the crucial role played by myeloid cell-derived XOR in LPS induced ALI *in vivo*. These data demonstrated that knockout of XOR in myeloid lineage cells *in vivo* reduced expression of XOR, modulated the development of ALI, reduced excretion of UA to the extracellular space, and modulated inflammatory activation in purified BAL macrophages from LPS induced ALI and in LPS stimulated BMDM and TGEM. While XOR was found to contribute broadly to macrophage inflammatory activation it also linked LPS induced lung inflammatory injury to NLRP3 inflammasome expression and metabolic state.

In the lung BAL of LPS insufflated mice, XOR was expressed in both newly recruited macrophages and in resident alveolar macrophages, suggesting a previously unrecognized role for XOR in the resident alveolar macrophages. A large fraction of XOR expressing adherent and interstitial mononuclear phagocytes was also observed that may contribute to LPS induced ALI as previously described for cytokine induced ALI [9–11]. The potential role of neutrophil specific XOR in the present analysis is unknown. In previous experiments, XOR could not be demonstrated in neutrophils derived from cytokine or LPS induced ALI *in vivo* nor induced by neutrophil differentiation *in vitro* [9, 10, 28]. However, approximately 6% of the total BAL cell XOR was found in neutrophils recovered from the BAL of LPS insufflated mice, and the contribution made by neutrophils to macrophage activation as well as differences between macrophage and neutrophil signaling was identified in both LPS induced ALI *in vivo* and in LPS stimulated macrophages [29]. Further analyses will be needed to determine the potential contribution of neutrophil specific XOR to LPS induced ALI/ARDS.

Microarray analysis of LPS induced ALI in C57BL/6 mice had previously shown the effect of LPS on a wide range of genes and had demonstrated the dynamic nature of the time course of LPS induced lung inflammation [23, 29–31]. Based on these results we analyzed lung tissue microarray 8 and 24 hrs after LPS insufflation where most genes regulated by LPS were anticipated to exhibit the greatest response using XOR^fl/fl^ and XOR^fl/fl//LysM-Cre^ mice. Myeloid specific XOR knockout was associated with the coordinate down-regulation of genes associated with inflammation and up-regulation of genes encoding mitochondrial OXPHOS (Tables S1-S4). Down-regulation of inflammasome genes NALP3, IL-1β, and IL18rap by XOR myeloid specific knockout 8 and 24 hrs after LPS exposure suggested an important function for XOR in mediating inflammasome expression..

The highly significant up-regulation of genes associated with OXPHOS and mitochondrial translation by myeloid specific XOR knockout in LPS insufflated mice was unexpected. These data revealed a role for wild-type XOR in modulating OXPHOS that may contribute to glycolysis in macrophages from the LPS treated lung. Inflammatory activation of macrophages with LPS has demonstrated the increased glycolytic state associated with LPS, whereas alternatively activated macrophages were found to be principally dependent on OXPHOS [32]. Data shown here identify a contribution to the glycolytic state in LPS insufflated mouse lungs and LPS stimulated BMDM that is mediated by XOR whereas knockout of macrophage XOR contributes to macrophage OXPHOS by up-regulation of *Tfb2m* and other genes involved in mitochondrial biogenesis.

The dual effect of XOR on NLRP3 inflammasome expression and OXPHOS is especially significant. Published results have shown the involvement of ROS, and mitochondrial ROS specifically, in activation of the NLRP3 inflammasome [27, 33, 34], and our results identify XOR as a physiologically significant source of ROS that promoted expression of the NLRP3 inflammasome and modulated mitochondrial ROS generation, perhaps reflecting the more robust expression of OXPHOS in the XOR knockout macrophages. Our data did not support a role for the NADPH oxidase as a source of ROS mediating NLRP3 and IL-1β expression. In this regard, knockout of the NADPH oxidase using p47*^phox-/-^* has revealed a role for NADPH oxidase derived ROS in limiting lung inflammation that produces an exaggerated response to LPS [35–37], consistent with results shown here that pharmacological inhibition of NADPH oxidase failed to block macrophage inflammatory activation.

A ‘two-step’ process for NLRP3 inflammasome-mediated IL-1β release is now widely accepted. The first step requires NF-kB mediated induction of IL-1β and NLRP3 expression, while the second step involves assembly of the NLRP3 inflammasome, caspase-1 activation, and then processing of pro-IL-1β and pro-IL-18 into mature cytokines that are then rapidly secreted [34, 38]. Nonetheless, genetic or prolonged pharmacological inhibition of IKKβ was found to enhance both IL-1β secretion and LPS injury *in vivo* [29]. While the potential contribution of XOR derived ROS to metabolic activation of the NLRP3 inflammasome was suggested previously [38], data shown here revealed that XOR derived ROS contribute not only to NF-κB mediated inflammatory gene expression, but also to expression of Nrf-2β which, although involved in anti-inflammatory gene expression, has also been recognized as critical mediator of proteolytic activation of the NLRP3 inflammasome and a source of ROS itself if chronically activated [27]. These data support a role for XOR derived ROS at the first step of IL-1β and NLRP3 expression, but also reveal a down-stream effect on IL-1β and IL-18 secretion through effect on redox status and redox response of the cells by fine tuning the balance between ROS and ROS derived antioxidants (step 2)[39] [40].

We observed significant differences in response to XOR ablation in BAL macrophages and BMDM that suggest differences in inflammasome activation by LPS in these different macrophage populations [24]. Ablation of XOR in BAL macrophages markedly reduced proteins for NLRP3, Pro- and mature-IL-1β, as well as Pro- and mature- caspase-1. Furthermore, secretion of both IL-1β and IL-18 proteins was markedly attenuated in BAL macrophages by XOR ablation. It is important to note that data shown here measured spontaneous IL-1β and IL-18 secretion in the absence of ATP or other inducers, and the level of spontaneously secreted proteins is approximately 100 fold lower than that reported for ATP or OCP induced secretion of LPS primed macrophages [41, 42]. In contrast, XOR ablation in BMDM failed to reduce Pro-IL-1β and reduced Pro-caspase-1 by only two fold. Furthermore, while secretion of mature IL-1β was dependent on XOR expression, it was independent of both NAC inhibited ROS or the NADPH Oxidase, suggesting a role for XOR in IL-1β secretion that was mediated by either macrophage metabolic state or intracellular UA as previously suggested [43]. These data are consistent with the defect identified in IL-1β secretion in M-CSF differentiated BMDM and monocyte derived macrophages that is not observed in circulating monocytes or BAL macrophages [41, 42].

Limitations of the present study include that they do not address whether the mechanism described here would apply to other stimuli, such as pathogen associated molecular pattern (PAMP). Furthermore, the model of acute lung injury used in this report is more focused on the initial disease process in respiratory failure and not the severe form of the injury that can be observed in conditions including sepsis. Further, we did not address whether XOR is necessary for the activation of other inflammasomes including AIM2 and NLRC4 and inflammasome assembly such as ASK. While our genetic model and pharmacologic XOR inhibitors used in this study show the improvement of metabolic readouts in the LPS treated macrophages in vitro and in vivo, in part through effects on mitochondrial electron transport chain (ETC), they don’t elucidate the differential ETC effects on NLRP3 inflammasome activation. Nevertheless, the genetic tools used in this study could be helpful in elucidating the potential interaction between XOR and different components of mitochondrial ETC.

In conclusion, this study has demonstrated the obligatory role of XOR in macrophage inflammatory activation that contributes to LPS induced ALI in mice. These data support a mechanism by which XOR derived ROS, acting in part through NF-κB, and Nrf-2β contribute broadly to inflammatory activation and specifically to expression and efficient activation of the NLRP3 inflammasome while at the same time contributing to macrophage glycolytic state by reducing expression of genes encoding mitochondrial OXPHOS via the mitochondrial transcription factor *Tfb2m* (Figure 7J). We suggest that XOR provides a physiologically important source of ROS that may act to promote macrophage inflammatory activation in balance with ROS induced antioxidant response that appear to play an important role in balanced redox remodeling ensuring efficient inflammasome activation and suggest that targeting XOR or molecules regulated by XOR may advance treatment of inflammatory disorders induced by prolonged or uncontrolled inflammasome activation.

## EXPERIMENTAL PROCEDURES

### Generation of XOR^fl/fl//LysM-Cre^ Mice

XOR^fl/fl//LysM-Cre^ mice were generated by crossing XOR^fl/fl^ mice with B6.129P2-*Lyz2^tm1(cre)Ifo^*/J (The Jackson Laboratory, strain # 004781). Mice were selected to be heterozygous for Cre recombinase and homozygous for the XOR “floxed’ allele (XOR^fl/fl^) as described [44, 45] and backcrossed into XOR^fl/fl^ six times. The generation of XOR knockout mice and derivative strains described in this study was approved by the University of Colorado Animal Care and Use Committee under the protocol number 49809(11)1E.

### Intratracheal administration of LPS

Experimental mice were treated with either saline or 5 mg/kg LPS. LPS (from Escherichia coli 055:B5) was dissolved in 0.9% NaCl, and mice were either treated with 100 μl LPS solution or saline (0.9% NaCl) using Intubation-mediated intratracheal (IMIT) administration. Briefly mice were sedated using isofluorine in Induction chamber for small animals. The mice were then put on mouse intubation stand and intubated using IV cannula, 22 gauge[46].

### Purification and Treatment of TGEM, BMDM, and Lung Macrophages

To generate purified Thiogylcollate Elicited Macrophages (TGEM), mice were injected intraperitoneally with 1 ml of 40mg/ml thioglycollate to elicit leukocyte infiltration. After 72 hrs peritoneal cells were harvested by saline flush and macrophages purified by adhesion to plastic dishes in RPMI 1640 with 10% FBS. Non-adherent cells were rinsed off after 1 hour. Adherent cells were transfected with Cre cDNA expression plasmid (Addgene, Cambridge, MA; #11916) or the empty vector using Lipofectamine according to the suppliers protocol (Invitrogen) (transfection efficiency of 70%). Adherent cells were photographed under light microscopy 24 hrs later and were quantitated by counting 20 fields per sample. Adherent cells were stained for XOR (red), nuclei (DAPI in blue), and cell shape actin (green), and representative high field photos are shown (Figure S1D). To generate Bone Marrow Derived Macrophages (BMDM), mice were killed by cervical dislocation. Tibia were dissected and flushed with cold medium into 50ml conical tubes. Cells were centrifuged for 5 min at 200 x g and resuspended 10 ml L-cell conditioned medium (LCCM). Cells were then cultured in LCCM at 1.0E6 cells/well in the presence of 25 ng/ml M-CSF for 6 days at 37°C under standard conditions as described [47]. XOR activity and protein level were determined after 6 days in culture. Alveolar macrophages, comprising both the resident macrophage population and the newly recruited macrophages, were obtained by lavage of lungs from mice insufflated with LPS and purified by adhesion to plastic culture dishes as described [9] or subjected to FACS sorting as described (Figure S2).

### Lung Tissue Nitrotyrosine Staining

Nitrotyrosine staining was performed largely as described previously [9]. Briefly, paraffin-embedded lung tissue sections were deparaffinized in ethanol and rehydrated in H2O and PBS. Slides were blocked with a solution of 7.5% normal goat serum, 2.5% β-casein, 0.1% triton X-100 for 3 h at room temperature. The primary antinitrotyrosine antibody Ab (Upstate Biotechnology, 06-284) was applied at a dilution of 1:1,000 for 15 min at room temperature in the blocking solution. Slides were washed in PBS, covered in Anti-Fade, and sealed under glass coverslips. 3-NT prebinding negative controls were performed by mixing the antinitrotyrosine antibody with 10 mM 3-nitrotyrosine in PBS for 1 h at room temperature before its addition to blocked slides exactly as described [9](Figure S1B).

### SOD/Catalse Activity

Superoxide dismutase (SOD) activity was measured using cytochrome c reduction inhibition as described [48]. Briefly, whole cell extracts were added to a mix of 50 mM xanthine, 10 mM cytochrome c, and 0.1 mM EDTA in 50 mM phosphate buffer pH 7.8. The reaction mix was homogenized and the rate of change of absorbance at 550-526 nm monitored with a diode array spectrophotometer at 25°C. SOD activity was calculated as described [48]. Catalase activity was determined as described using fresh stock concentrations of H2O2 [49]. Whole cell extracts were added to 20 mM H2O2 in 50 mM potassium phosphate pH 7.0 and incubated at 25°C, recording absorbance changes at 240 nm. Activity was determined using an extinction coefficient of 43.6 M-1.

### XOR Activity

XOR activity was determined at 293 nm by quantitation of Allopurinol-inhibited uric acid synthesis using a Beckman-Coulter DU640 spectrometer, 1-cm path length, and a molar extinction coefficient for uric acid of 12,600 M_1. All XOR activity data show nanomoles of uric acid/min/mg of total protein. D-form XOR activity was determined in the presence of NAD+ while O-form was determined in its absence [9].

### Mitochondrial ROS Analysis

BAL cells were harvested from LPS or saline insufflated mice and stained with MitoSOX Red indicator (Invitrogen) for flow cytometric detection of mitochondrial superoxide in the presence of either Alexaflour conjugated anti- F4/80 or anti-Ly6G as described [50]. Briefly, cells were incubated in DMEM, 10% FBS at 37°C with 25 nM MitoTracker Green for 15 min in the dark followed by addition of 4 μM MitoSox Red for a further 15 min. Stained cells were washed twice in PBS and analyzed by flow cytometry or Zeiss LSM 700 confocal microscopy. Mean fluorescence intensity (MFI) from Mitosox was determined on flow analyzed cells, and data were analyzed by one sided ANOVA representing the MFI for six independent measurements (**, p < 0.02).

### Assessment of mitochondrial membrane potential

We used tetramethylrhodamine ethyl ester (TMRE) staining assay kit to assess mitochondrial membrane potential. We followed the method of Dingley, et al [51] and also the provided instruction by the company for TMRE analysis. Briefly cells were incubated with TMRE for 15-30 min, then were washed with PBS / 0.2% BSA. The stained pellet/adherent cells were analyzed with flow cytometer using 488nm laser for excitation and at emission 575 nm, or fluorescent microscope. We used FCCP (carbonyl cyanide 4-(trifluoromethoxy) phenylhydrazone), which is a ionophore uncoupler of oxidative phosphorylation for our positive control.

### Analysis of mitochondrial respiration using Clark electrode

For oxygen consumption measurements, cells were trypsinized, centrifuged at 1200 x g for 10 min, resuspended in 500 ul of medium containing 20 mM HEPES, 10 mM MgCl2, 250 mM sucrose (pH 7.3), and counted. Aliquots of 1 × 106 cells were combined with 500 ul of the respiration medium containing 130 mM KCl, 20 mM HEPES, 5 mM KH2PO4, 2 mM MgCl2, 0.1 mM EGTA, and 10 μg/ml digitonin and mixed for 10 min. Optimal concentration of digitonin was determined in concentration-dependent experiments, in which digitonin was tested in the range of 2.5-20 ug/ml. The respiration rate of mitochondria in the permeabilized cells was measured with Clark-type oxygene electrode (Oxygene electrode Units DW1; Hansatech Instruments, Norfolk, UK) at 37 ^0^C. After the basal respiration rate was stabilized, oxygen consumption measurements were initiated by the addition of complex I substrates, 2 mM pyruvate, 3.2 malate and 20 mM glutamate followed by the addition of 1 mM ADP to initiate a state 3 respiration at complex I. After oxygen consumption reached saturation, 1 uM rotenone (inhibitor of complex I), 2 mM succinate (complex II substrate) and 1 mM ADP were added to initiate the basal and state 3 respiration at complex II, respectively. The respiration was terminated by the addition of Oligomycin (4 ug/ml). Overall, O2 consumption was monitored for about 15 min.

### IKK Kinase Assay

Kinase assays were performed as previously described by our group [52]. Control 293T cells and macrophage cells were lysed and immunoprecipitated with anti-IKKγ antibody (BD)(Figure S5B) . The immune complexes were incubated with 1 μg GST-IκBα(1–54) in the presence of 10 μCi γ[32P]ATP at 30 °C for 30 min. Proteins were resolved by SDS-PAGE and transferred to a nitrocellulose membrane that was exposed to x-ray film.

### Microarray Analysis

Eight or 24 hrs after LPS insufflation lungs were perfused blood free, dissected, and snap frozen in liquid nitrogen. After pulverization over liquid nitrogen, total lung RNA was prepared using Tri-Reagent (Sigma) extraction. Total RNA was washed in 70% ethanol, quality checked by gel analysis, and transferred to the UCD Microarray Core laboratory. cDNA generation, probe preparation, and annealing reactions, were performed by the core laboratory using Affymetrix Mouse Specific Gene 2.0 ST array and Sample Labeling Reagents for Affymetrix Exon/Gene 1.0/2.0 Arrays. Log2 normalized and annotated data were prepared on Excel worksheets for each chip, and comparison between groups was conducted by determining ratios of either +LPS/-LPS or XOR^fl/fl//LysM-Cre^/XOR^fl/fl^ when both groups were treated with LPS. Data were scored for up or down regulation and data showing >1.5 fold regulation were collected and analyzed using the DAVID version 6.7 platform [53, 54] and verified by the Array Express open portal (http://www.ebi.ac.uk/services). This produced sets of 564 genes down-regulated and 214 genes up- regulated at 24 hrs by XOR knockout. Gene identifications were made using GENBANK_ACCESSION number, REFSEQ_mRNA, ENSEMBL_TRANSCRIPT, ENTREZ_GENE-ID, and the OFFICIAL_GENE_SYMBOL. Up or down regulated genes were sorted first by p-Value using Fishers Exact Test Score and the Bonferonni correction for false discovery as described [53, 54] and subsequently sorted for enrichment score. Finally, data sets were sorted by Functional Annotation Clustering [53, 54] and functional groups showing p<0.0076 have been tabulated (Tables S1-S4).

### Fluorescence Lifetime Imaging (FILM) and Phasor Plot Analysis

Fluorescence intensity and lifetime imaging of two-photon excited NADH was performed at the UCD bioimaging core laboratory largely as described [55, 56]. The lifetime measurements were carried out on a Zeiss LSM780 (Carl Zeiss, Jena, Germany) confocal microscope equipped with ISS FastFLIM technology and a titanium:sapphire Chameleon Ultra II (Coherent, Santa Clara, CA). Briefly, the intrinsic NADH fluorescence was observed through a 40x water immersion objective Zeiss Korr C-Apochromat NA 1.2 (Carl Zeiss, Jena, Germany) for monolayer cell culture. The NADH fluorescence was isolated using a 488 nm long pass dichroic mirror (Semrock Di02-R488-25-D) and a bandpass filter the bandpass filter with the spectral range 415-485 nm (Semrock FF01-450/70-25) and detected with a Hamamatsu H7422p-40 photon-counting PMT connected to a ISS A320 FastFLIM box (ISS, Champaign, IL). Lifetime calibration was done using fluorescein aqueous solution of 30 nM for which lifetime is known (4 ns). Calibration and NADH imaging of the live cells were done using the same acquisition parameters. S and G parameters are polar coordinates for the lifetime decay in the phasor plot representation, and detailed discussion of their derivation have been published [56, 57]. In the current data, phasors for a single decay (bound 3.2 ns, free 0.4 ns) are located on the universal circle such that a phasor corresponding to the short halflife is close to the point (1,0) while a phasor corresponding to a very long lifetime will be close to the point (0,0). Importantly, the ratio of free and bound NADH lie on a straight line that readily enables determination of bound/free ratio as described [56] when XOR^fl/fl^ and XOR^fl/fl//LysM-Cre^ data are presented on the same phasor plot as shown in Figure 5.

### Cell Viability and Survival

Viability and survival of macrophages was determined either by sulforhodamine-B (SRB) assay exactly as described [58] for cells plated at 1.0E6 cells/well or by trypan blue exclusion assay as described [11].

### Real Time Quantitative RT-PCR

qRT-PCR was performed on total RNA and normalized/shown as relative mRNA expression to HPRT as described previously [10]. Primer sequences used for qT-PCR are shown in Table S5. Primers not listed (CD36, CD206, and MCP-1) were purchased prepackaged from Life Technologies (Grand Island, NY, USA) or Amersham. Reactions were performed using the TaqMan kit (LifeTechnologies) as specified.

### ELISA and Western Blot

Cytokine protein levels were determined either by ELISA (Enzyme-Linked ImmunoSorbent Assay) using kits from R&D Systems Inc. or by western blot as described previously after loading 50 ug protein/well [10]. ELISA data were quantitated on a Luminex 100 reader according to the manufacturer’s instructions. Antibodies and sources used in the present analysis have been annotated in Table S6 and were routinely diluted 1:1000 for western blot analysis.

### Statistical Analyses

Data are expressed as the mean and standard error of the mean and were assessed for significance using either the Student’s t-test or ANOVA. A *p* value of <0.05 was considered significant. Statistical analysis of microarray data was performed as described above using the DAVID platform.

## SUPPLEMENTAL INFORMATION

Supplemental Information includes select methods, data figures, and tables.

## AUTHOR CONTRIBUTIONS

R.M.W. and M.A.F. conceived and designed the experimental protocols. M.A.F., J.M., M.L., E.G., P.P., K.W., M.G.F., S.C.P., performed the experimental analyses. All authors contributed to analysis of the data. D.B., W.J., M.K., Y.R.U ., D.I and K.R.S., contributed to the critical evaluation of the manuscript. R.M.W. and M.A.F. wrote the manuscript.

## ACKNOWLEDGEMENTS

The authors would like to thank, Dr. Radu Moldovan, Dr. Daniel Hernandez, Dr. Suzette R. Riddle, Dr. Jerry Wong, Dr. Ruby F. Fernandez-Boyanapalli and Dr. Richard Scarpulla for expert technical assistance. The authors thank the National Institutes of Health (HL113809, RMW; HL152961, KS), The American Thoracic Society (MAF), the Department of Defense (W81XWH-14-1-0451, MAF), and University of Colorado CCTSI (KL2TR001080, MAF) for support of this work.

## SUPPLEMENTAL FIGURE LEGENDS

**Figure S1.**
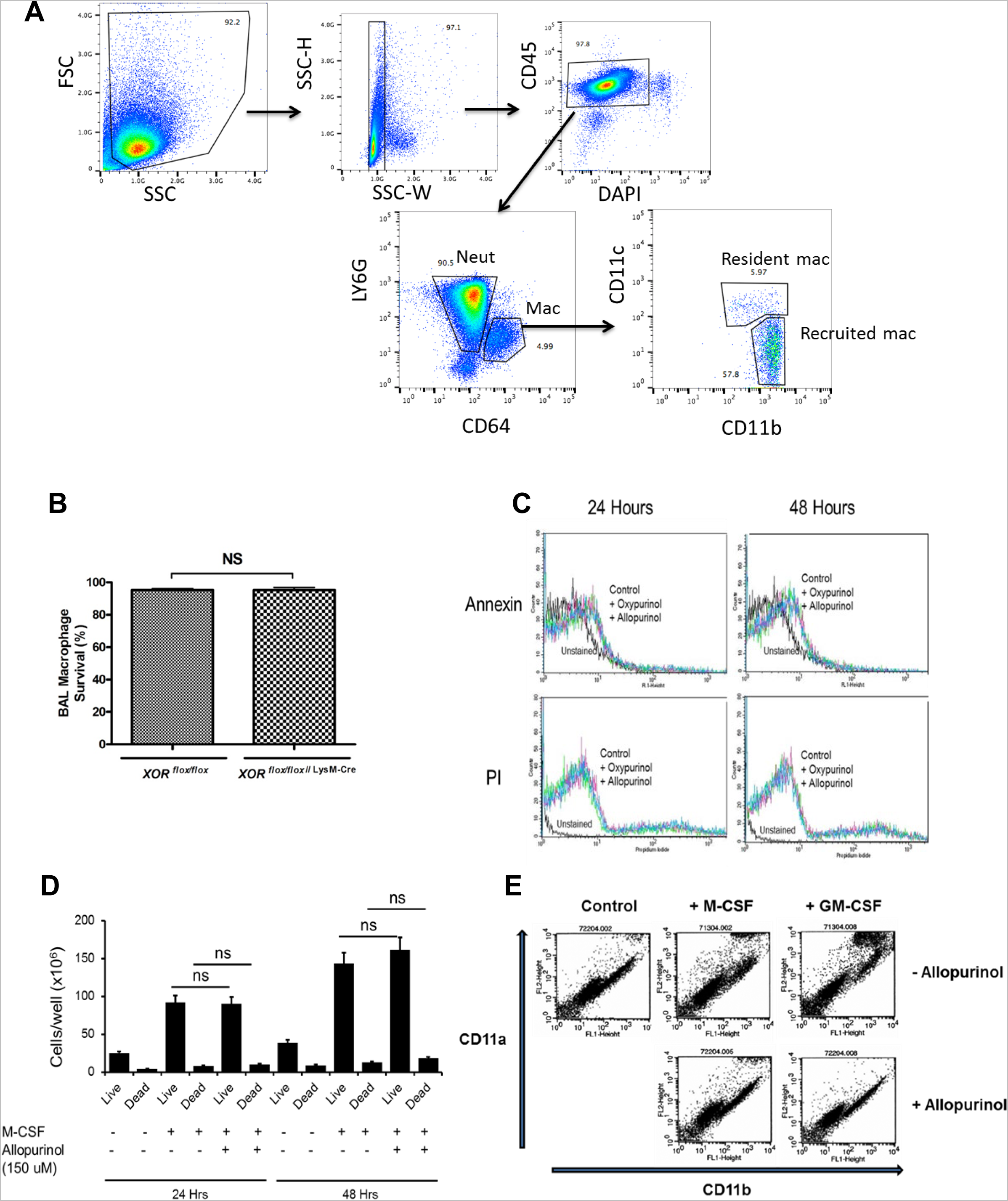
Flow cytometry gating strategy for MNP/PMN sorting. (A) Gating strategy used for isolation of neutrophils, resident macrophages, and recruited macrophages from mouse bronchoalveolar fluid is shown. Briefly, recruited macrophages, resident macrophages and neutrophils were isolated using flow cytometry cell sorting from BAL fluid obtained from 5 pooled control mice using an MoFlo XDP-100 flow cytometer (Beckman Coulter). Monoclonal antibodies (mAb) were purchased from BD biosystems: Allophycocyanin-conjugated mAb to CD64; allophycocyaninCy7-conjugated mAb to CD45; PE-CF594-conjugated mAb to Ly6G; FITC-conjugated mAb to CD11c; BV510-conjugated mAB; EF450 conjugated mAb’s to B220, CD3. Dead cells were excluded using DAPI (4’,6-diamidino-2-phenylindole). Recruited macrophages were identified as CD45+, CD64+, CD11b+, CD11c^lo^. Resident macrophages were identified as CD45+, CD64+, CD11c+, CD11b^lo^ . Neutrophils were identified as CD45+, Ly6G+, CD64-. Purity was improved by excluding dead cells with DAPI, singlets with SSC-W, T cells with CD3 and B cells with B220. Cells were sorted directly into Hanks Balanced Salt Solution media (Life Technologies) supplemented with 1% Hyclone fetal bovine serum (Thermo Scientific). 1.0 E6 cells of each fraction were collected for analysis of XOR activity. Neut (neutrophil), Mac (macrophage). (B) SRB uptake in macrophages (resident and alveolar) purified from the lungs of XOR^fl/fl^ and XOR^fl/fl//LysM-Cre^ mice 48 hrs after LPS insufflation. Data show % SRB negative cells; n=3; ns, not significant. (C) Purified macrophages (resident and alveolar) were allowed to adhere for 1 hr on plastic culture dishes, washed, and then treated with oxypurinol (150 uM) or allopurinol (150 uM). After 24 or 48 hrs cells were stained with antisera to annexin V and propidium iodide (PI). Annexin V and PI were detected using the Annexin V Apoptosis Detection Kit APC (Affymetrix eBioscience Catalog Number: 88-8007), and cells subjected to FACS analysis. Signals were color coded as follows: Unstained cells (black), untreated controls (yellow), oxypurinol treated (blue), allopurinol treated (purple). (D) Purified macrophages were prepared as in panel B and treated with allopurinol (150 uM) *in vitro* in the presence or absence of 10.0 ng/ml M-CSF. After 24 or 48 hrs cells were stained with trypan blue, counted, and scored as live (no stain) or dead (blue) as described (Parmley et al., 2007). Data were analyzed by one sided ANOVA. ns refers to intra-group comparison between cells treated with M-CSF alone or in combination with allopurinol. (E) Purified macrophages were treated with allopurinol (150 uM) *in vitro* in the presence or absence of 10.0 ng/ml M-CSF or GM-CSF. After 24 hrs cells were stained for surface expression of CD11a and CD11b, subjected to FACS analysis, and scatter diagrams obtained. Both M-CSF and GM-CSF treatment resulted in the production of a CD11a*^Hi^*/CD11b*^Hi^* population that was blocked by Allopurinol inhibition.

**Figure S2.**
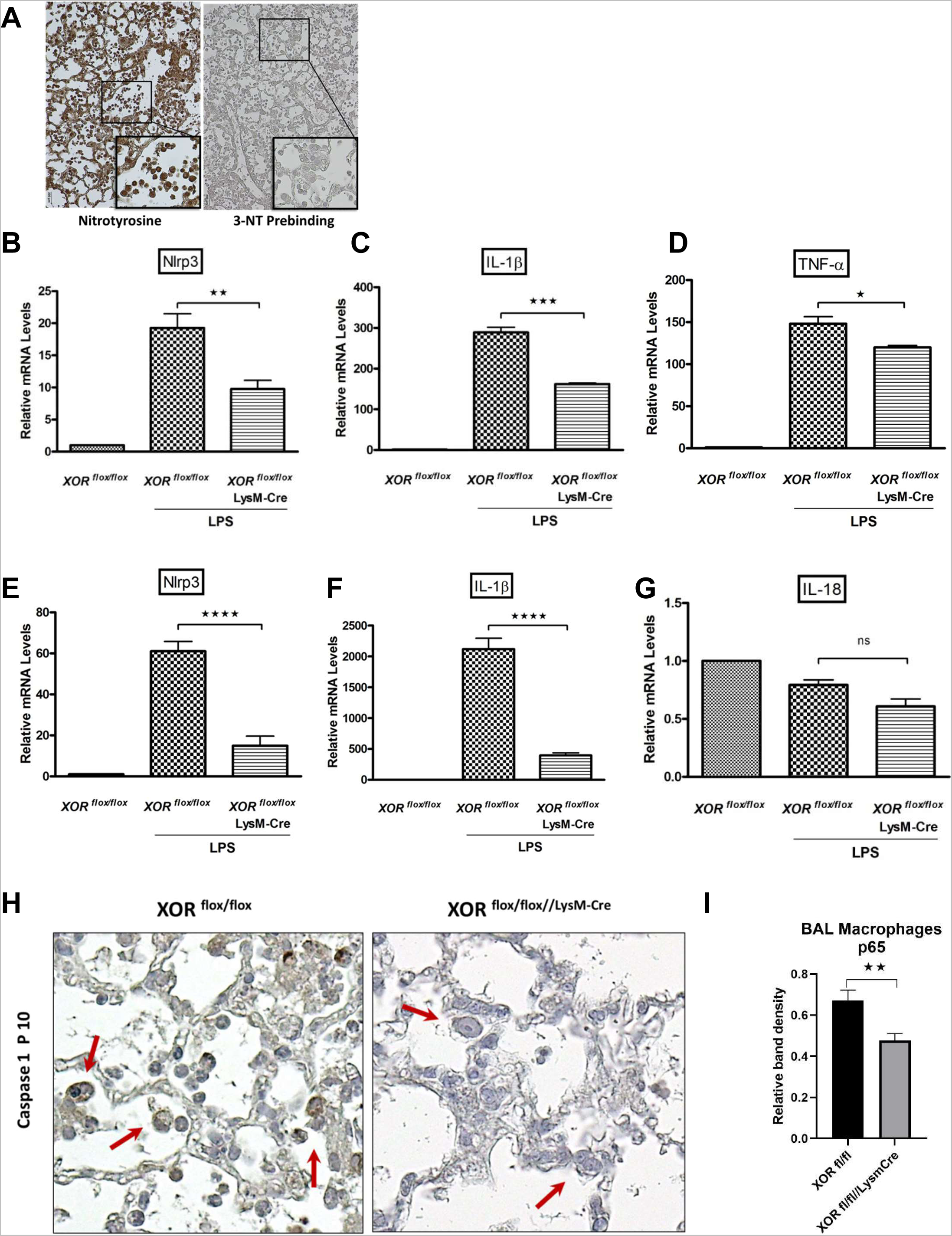
Lung Tissue and macrophage inflammatory activation following LPS insufflation. (A) Paraffin embedded lung tissue specimens were stained for the presence of nitrotyrosine. Representative photos show both alveolar macrophage staining (top) and perivascular macrophage staining (bottom). 3-NT prebinding control was performed on lung tissue slides from XORfl/fl mice using 10 mM 3-NT (Sigma) that was added to antibody for 60 min on ice prior to staining the slides. (B-D) Quantitative RT-PCR analysis of whole lung for NLRP3, IL-1β, and IL-18. RNA isolated 48 hrs after saline or LPS insufflation in XOR^fl/fl^ and XOR^fl/fl//LysM-Cre^ mice. Data show mean and S.E. of six independent measurements for each group, and data were analyzed by Student’s t-test. (*, p<0.05; **, p<0.02; ***, p<0.001). (E-G) RNA was obtained from purified macrophages from saline or LPS insufflated XOR^fl/fl^ and XOR^fl/fl//LysM-Cre^ mice, and quantitative RT-PCR was performed for NLRP3, IL-1β, and IL-18. Data show the mean and S.E. of six mice in each group and were analyzed by Student’s t-test (****, p<0.0005; ns, not significant). (H) Lung tissue histological slides from XOR^fl/fl^ and XOR^fl/fl//LysM-Cre^ mice insufflated with LPS 48 hrs before were stained with HRP conjugated antisera against caspase-1 p10 protein. (I) Relative band density of whole cell lysates of BAL macrophages purified from lungs of XORfl/fl and XORfl/fl//LysM-Cre mice insufflated with LPS 48 hrs before and probed with antisera against NFκB p65.

**Figure S3.**
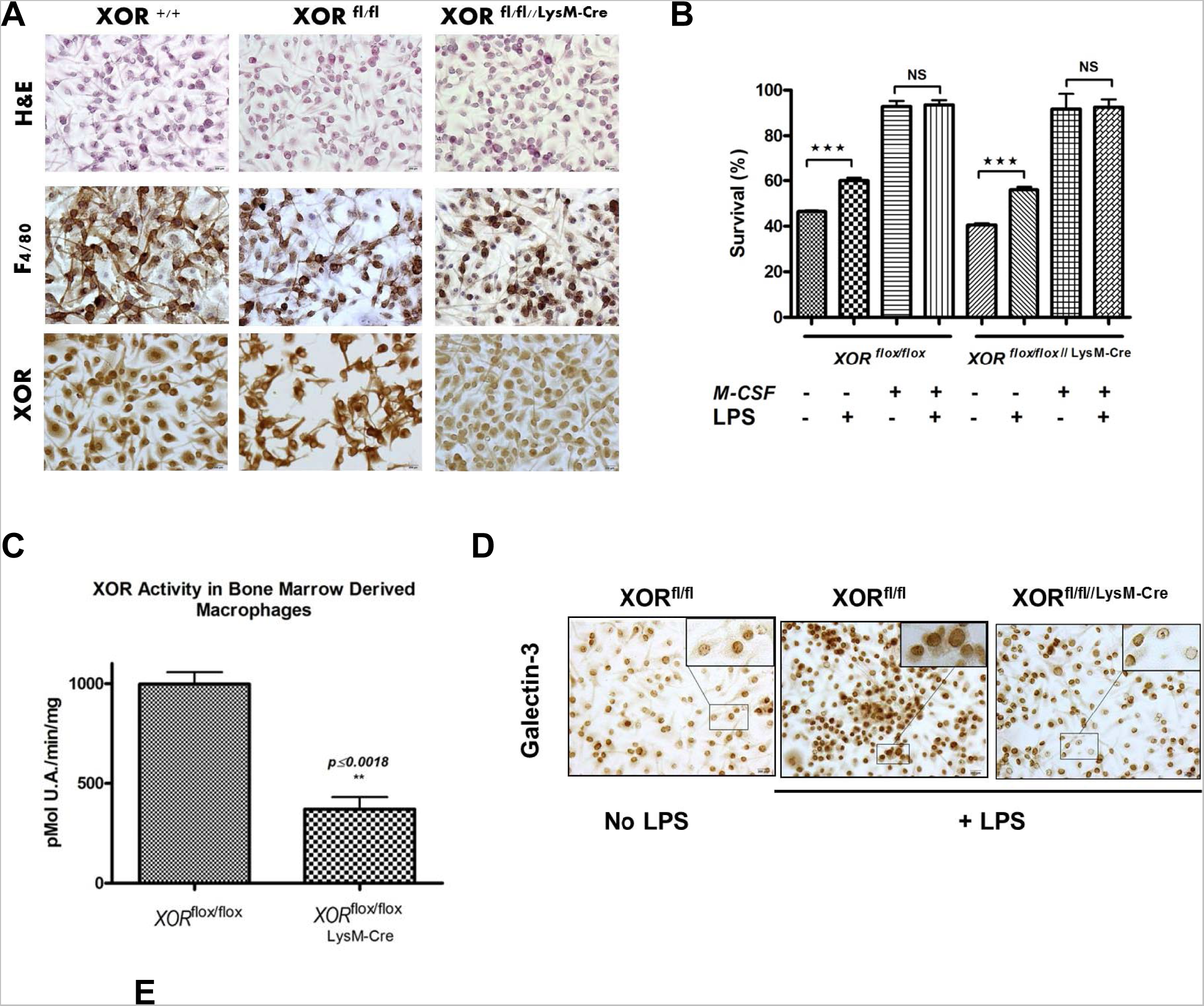
Myeloid Specific XOR Knockout *In vivo* Reduces Inflammatory Activation of BMDM. (A) BMDM were generated from XOR+/+ (C57BL/6), XOR^fl/fl^, and XOR^fl/fl//LysM-Cre^ mice and purified by adhesion to plastic culture dishes. After six days, cells were harvested, stained by H&E or for the macrophage specific marker F4/80, and XOR antigen. (B) Cell number and survival were determined by SRB assay. (C) XOR activity was determined from six independent BMDM preparations and were analyzed by Student’s t-Test. Data show the mean and SE. (D) Mouse cytokine ELISA array (Signosis, Inc., Santa Clara, CA, USA) was used on whole cell lysates from these cells as described by the supplier. Data were normalized to the untreated WT control and shown as the mean and S.E. XORfl/fl and XORfl/fl//LysM-Cre data were analyzed by Students t-test (*, p<0.05; **, p<0.02).

**Figure S4.**
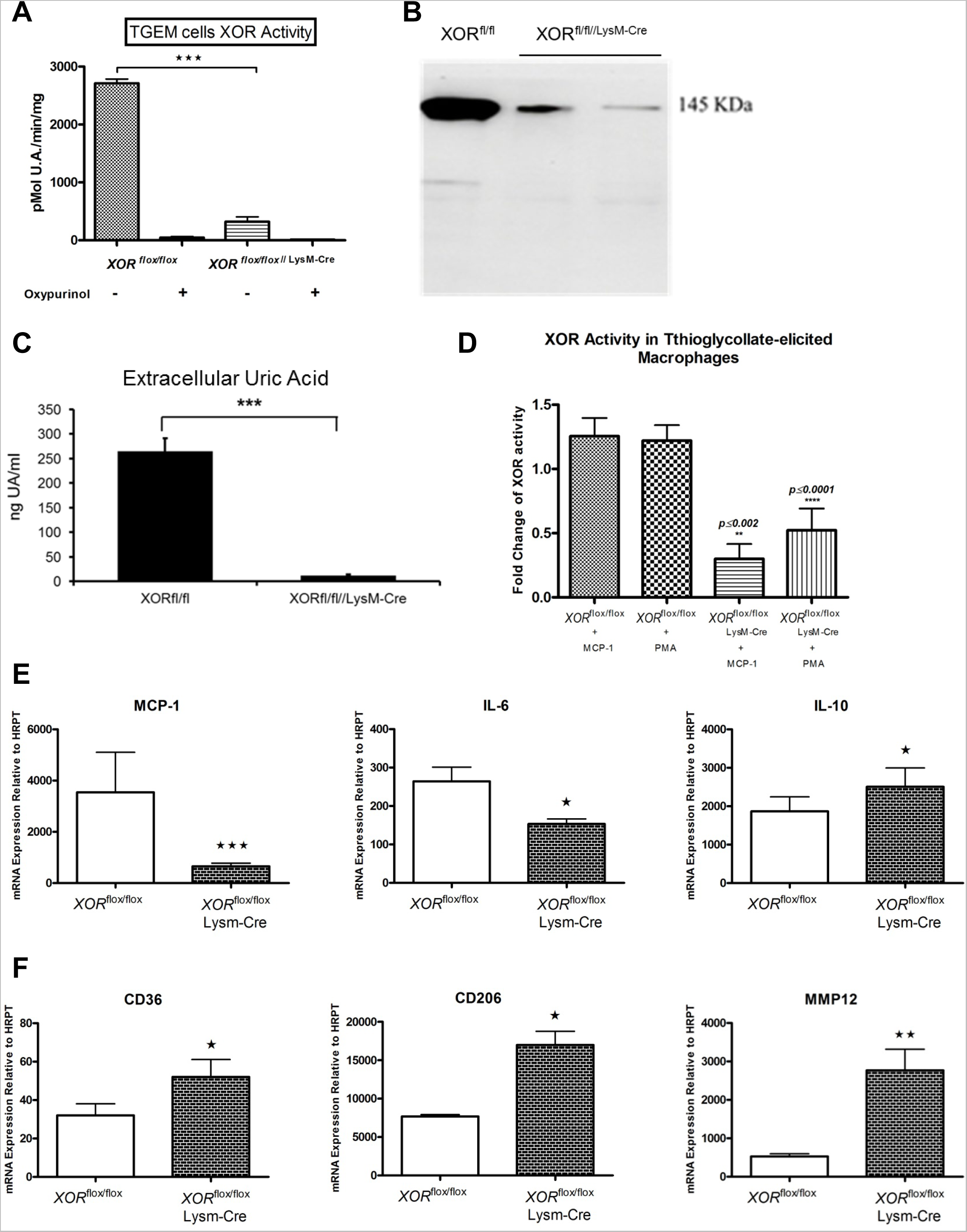
Myeloid Specific XOR Knockout *In vivo* Modulates TGEM Inflammatory State. (A) TGEM were produced from XOR^fl/fl^ and XOR^fl/fl//LysM-Cre^ mice as indicated in Figure 1, purified by adhesion to plastic culture dishes, and assayed for XOR activity after 24 hrs of *in vitro* culture. Data show the mean and SE of six independent measurements. Oxypurinol was included in parallel assays to confirm XOR activity. (B) XOR western blot was run using one randomly selected XOR^fl/fl^ littermate control and two randomly selected _XOR_fl/fl//LysM-Cre _littermates._ (C) Excreted UA was measured in the cell free medium from the same specimens used in panel A. (D) XOR induction by MCP-1 and PMA is refractory in TGEM from XOR^fl/fl//LysM-Cre^ mice. TGEM were purified by adhesion to plastic culture dishes and after one hour were treated with either MCP-1 (10 ng/ml) or PMA (33 nM). Cells were harvested after 24 hrs and XOR activity determined as above. Data were normalized to the untreated XOR^fl/fl^ value. Data show the mean and SE of six independent measurements and were analyzed by one sided ANOVA. (E, F) TGEM were purified from six different XOR^fl/fl^ and XOR^fl/fl//LysM-Cre^ mice and placed into tissue culture for 24 hrs. Total cellular RNA was isolated and transcripts for different inflammatory genes (E) or markers (F) were measured by quantitative real time RT-PCR. Data were analyzed by Student’s t-Test.

**Figure S5.**
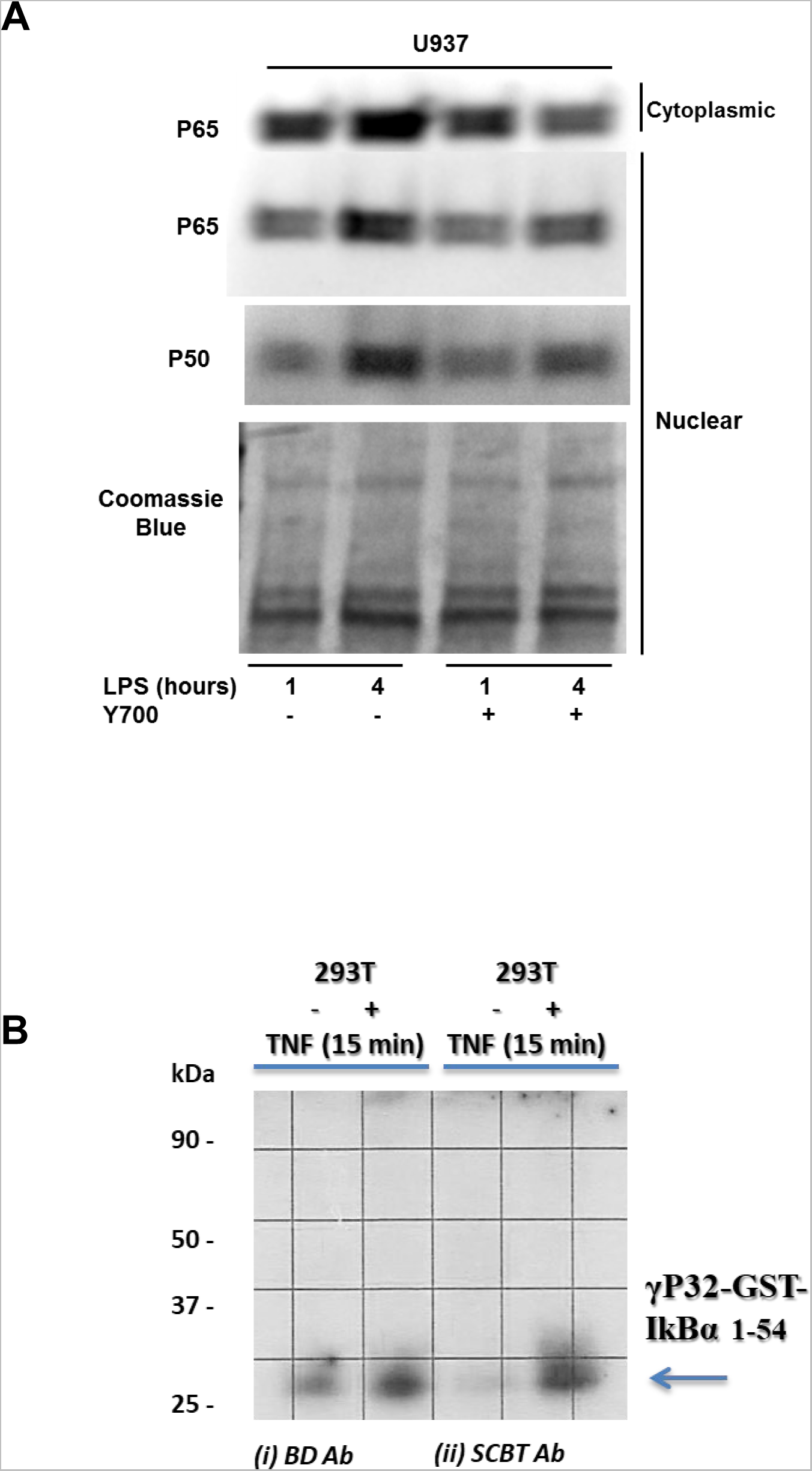
XOR pharmacological inhibition reduced nuclear NFκB accumulation in LPS treated human U937 cells. (A) U937 cells were grown *in vitro* and stimulated with LPS (100 ng/ml) for 1 or 4 hours in the presence or absence of the XOR inhibitor Y700 (50 nM). Y700 was used in this experiment instead of Allopurinol, because it lacks any known ROS scavenging capacity. Nuclear and cytoplasmic lysates were generated as described (Roberts et al., 2007; Seymour et al., 2006) and proteins subjected to western blot analysis of NFκB p65 and p50. Western blot is representative of three independent experiments. Coomassie Blue staining was used to control for loading of nuclear fractions. (B) Kinase assays in control 293T cells. cells were lysed and immunoprecipitated with anti-IKKγ antibodies y (BD and SCBT). The immune complexes were incubated with 1 μg GST-IκBα(1–54) in the presence of 10 μCi γ[32P]ATP at 30 °C for 30 min. Proteins were resolved by SDS-PAGE and transferred to a nitrocellulose membrane that was exposed to x-ray film.

**Figure S6.**
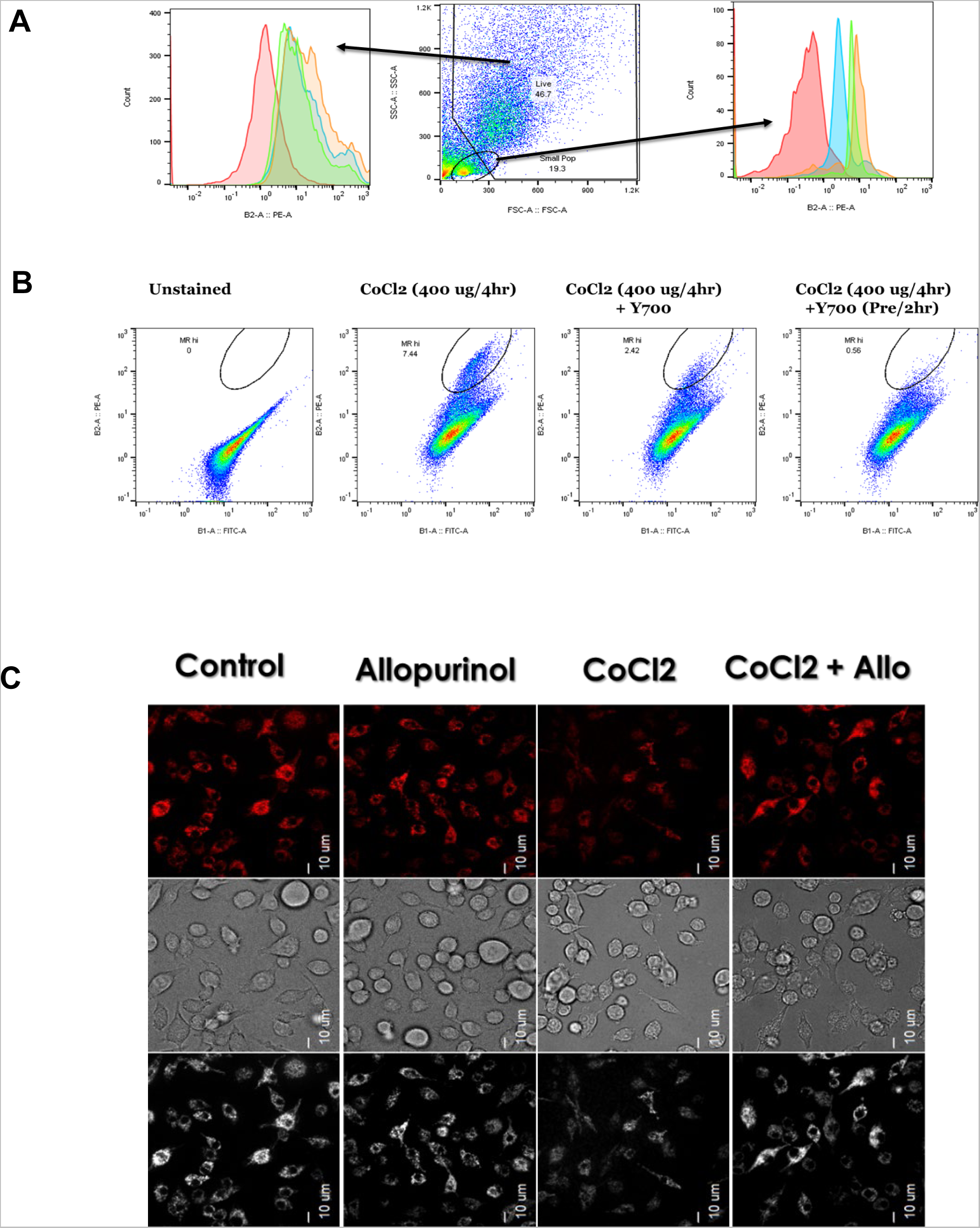
XOR pharmacologic inhibition attenuates LPS and CoCL2 mitochondrial dysfunction in BMDM and RAW246.7 macrophage cell line. (A) FACS analysis of BMDM cells stained with MitoSOX Red treated with LPS with and without XOR inhibitor Y700. (B) FACS analysis of RAW246.7 cells stained with MitoSOX Red treated with CoCL2 with and without XOR inhibitor Y700. The XOR inhibition reduces the Mitosox^high^ population in CoCL2 treated cells. (C) Mitochondrial membrane potential assessment using fluorescent microscopy in RAW246.7 cells stained with TMRE and treated with CoCL2 with and without Allopurinol.

## SUPPLEMENTAL TABLES

**TABLE S1.**
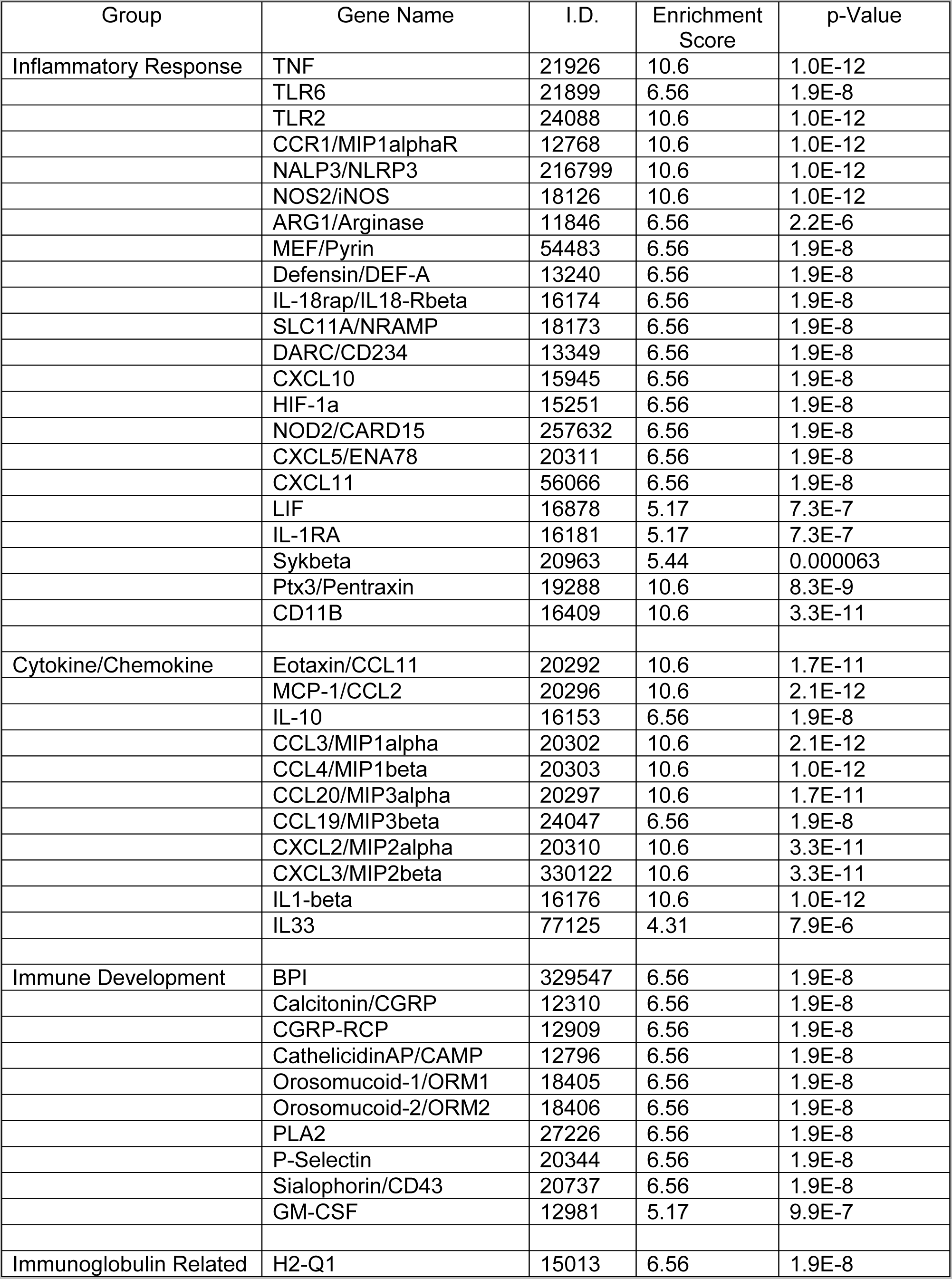
Summary of microarray data for genes down-regulated 8 hrs after LPS insufflation.

Genes Down-Regulated at 8 Hours
| Group | Gene Name | I.D. | Enrichment Score | p-Value |
| --- | --- | --- | --- | --- |
| Inflammatory Response | TNF | 21926 | 10.6 | 1.0E-12 |
|  | TLR6 | 21899 | 6.56 | 1.9E-8 |
|  | TLR2 | 24088 | 10.6 | 1.0E-12 |
|  | CCR1/MIP1alphaR | 12768 | 10.6 | 1.0E-12 |
|  | NALP3/NLRP3 | 216799 | 10.6 | 1.0E-12 |
|  | NOS2/iNOS | 18126 | 10.6 | 1.0E-12 |
|  | ARG1/Arginase | 11846 | 6.56 | 2.2E-6 |
|  | MEF/Pyrin | 54483 | 6.56 | 1.9E-8 |
|  | Defensin/DEF-A | 13240 | 6.56 | 1.9E-8 |
|  | IL-18rap/IL18-Rbeta | 16174 | 6.56 | 1.9E-8 |
|  | SLC11A/NRAMP | 18173 | 6.56 | 1.9E-8 |
|  | DARC/CD234 | 13349 | 6.56 | 1.9E-8 |
|  | CXCL10 | 15945 | 6.56 | 1.9E-8 |
|  | HIF-1a | 15251 | 6.56 | 1.9E-8 |
|  | NOD2/CARD15 | 257632 | 6.56 | 1.9E-8 |
|  | CXCL5/ENA78 | 20311 | 6.56 | 1.9E-8 |
|  | CXCL11 | 56066 | 6.56 | 1.9E-8 |
|  | LIF | 16878 | 5.17 | 7.3E-7 |
|  | IL-1RA | 16181 | 5.17 | 7.3E-7 |
|  | Sykbeta | 20963 | 5.44 | 0.000063 |
|  | Ptx3/Pentraxin | 19288 | 10.6 | 8.3E-9 |
|  | CD11B | 16409 | 10.6 | 3.3E-11 |
| Cytokine/Chemokine | Eotaxin/CCL11 | 20292 | 10.6 | 1.7E-11 |
|  | MCP-1/CCL2 | 20296 | 10.6 | 2.1E-12 |
|  | IL-10 | 16153 | 6.56 | 1.9E-8 |
|  | CCL3/MIP1alpha | 20302 | 10.6 | 2.1E-12 |
|  | CCL4/MIP1beta | 20303 | 10.6 | 1.0E-12 |
|  | CCL20/MIP3alpha | 20297 | 10.6 | 1.7E-11 |
|  | CCL19/MIP3beta | 24047 | 6.56 | 1.9E-8 |
|  | CXCL2/MIP2alpha | 20310 | 10.6 | 3.3E-11 |
|  | CXCL3/MIP2beta | 330122 | 10.6 | 3.3E-11 |
|  | IL1-beta | 16176 | 10.6 | 1.0E-12 |
|  | IL33 | 77125 | 4.31 | 7.9E-6 |
| Immune Development | BPI | 329547 | 6.56 | 1.9E-8 |
|  | Calcitonin/CGRP | 12310 | 6.56 | 1.9E-8 |
|  | CGRP-RCP | 12909 | 6.56 | 1.9E-8 |
|  | CathelicidinAP/CAMP | 12796 | 6.56 | 1.9E-8 |
|  | Orosomucoid-1/ORM1 | 18405 | 6.56 | 1.9E-8 |
|  | Orosomucoid-2/ORM2 | 18406 | 6.56 | 1.9E-8 |
|  | PLA2 | 27226 | 6.56 | 1.9E-8 |
|  | P-Selectin | 20344 | 6.56 | 1.9E-8 |
|  | Sialophorin/CD43 | 20737 | 6.56 | 1.9E-8 |
|  | GM-CSF | 12981 | 5.17 | 9.9E-7 |
| Immunoglobulin Related | H2-Q1 | 15013 | 6.56 | 1.9E-8 |

**TABLE S2.** Summary of microarray data for genes up-regulated 8 hrs after LPS insufflation.

Genes Up-Regulated at 8 Hours
| Group | Gene Name | I.D. | Enrichment Score | p-Value |
| --- | --- | --- | --- | --- |
| Oxidative Phosphorylation | ND1/NADH-Ubiquinone Dehydrogenase | 17716 | 3.08 | 2.9E-5 |
|  | ND2/NADH-Ubiquinone Dehydrogenase | 17717 | 3.08 | 2.9E-5 |
|  | ND3/NADH-Ubiquinone Dehydrogenase | 17718 | 3.08 | 8.0E-5 |
|  | ND4/NADH-Ubiquinone Dehydrogenase | 17719 | 3.08 | 8.0E-5 |
|  | ND5/NADH-Ubiquinone Dehydrogenase | 17721 | 3.08 | 3.0E-5 |
|  | ND6/NADH-Ubiquinone Dehydrogenase | 17722 | 3.08 | 3.0E-5 |
|  | CYTB/Cytochrome b | 17711 | 3.08 | 2.9E-5 |
|  | COX1 | 17708 | 3.08 | 2.9E-5 |
|  | COX2 | 17709 | 3.08 | 2.9E-5 |
|  | COX3 | 17710 | 3.08 | 2.9E-5 |
|  | COX 7A1 | 12865 | 3.08 | 2.9E-5 |
|  | ATPEpsilon | 67126 | 3.08 | 2.9E-5 |
|  | ATP6 | 17705 | 3.08 | 2.9E-5 |
|  | ATP8 | 17706 | 3.08 | 2.9E-5 |
| Mitochondrial Function | UCP2 | 22228 | 3.08 | 2.9E-5 |
|  | TST/mitochondrial thiosulfate transferase | 22117 | 3.08 | 2.9E-5 |
|  | CAT1/Crat | 12908 | 3.08 | 8.0E-5 |
|  | MRPL32 | 75398 | 18.05 | 6.3E-7 |
|  | MRPL41 | 670250 | 18.05 | 6.3E-7 |
|  | TufM | 233870 | 18.05 | 6.3E-7 |
|  | CysRS | 71941 | 18.05 | 6.3E-7 |
|  | 22 Trn Genes/tRNA Translation Elongation | 17726-17747 | 18.05 | 1.3E-25 |
|  | Rnr1/Ribosomal RNA precursor | 17724 | 18.05 | 6.3E-7 |
|  | Rnr2/Ribosomal RNA precursor | 17725 | 18.05 | 6.3E-7 |
| Cell Surface/Signaling | CD59a | 12509 | 3.94 | 4.7E-6 |
|  | CD40 | 21939 | 3.94 | 8.0E-5 |
|  | Adora1/ARA1 | 11539 | 3.94 | 4.7E-6 |
|  | CD95/Fas | 14102 | 3.94 | 4.7E-6 |

**TABLE S3.** Summary of microarray data for genes down-regulated 24 hrs after LPS insufflation.

Genes Down-Regulated at 24 Hours
| Group | Gene Name | I.D. | Enrichment Score | p-Value |
| --- | --- | --- | --- | --- |
| Inflammatory Response | TNF-R2 | 21938 | 3.13 | 0.000040 |
|  | TLR6 | 21899 | 3.13 | 0.000040 |
|  | TLR9 | 81897 | 3.13 | 0.000040 |
|  | IL1-beta | 16176 | 3.13 | 0.000040 |
|  | IL1-R2 | 16178 | 3.13 | 0.00038 |
|  | CCR1/MIP1alphaR | 12768 | 3.13 | 0.000075 |
|  | CCL4/MIP1beta | 20303 | 3.13 | 0.000040 |
|  | NALP3/NLRP3 | 216799 | 3.13 | 0.000040 |
|  | CXCR4 | 12767 | 2.74 | 0.000050 |
|  | NOS2/iNOS | 18126 | 3.13 | 0.00017 |
|  | ARG1/Arginase | 11846 | 3.13 | 0.00059 |
|  | ALOX5/LOX5 | 11689 | 3.13 | 0.00017 |
|  | BAX/BCL2-L4 | 12028 | 3.13 | 0.000075 |
|  | MEF/Pyrin | 54483 | 3.13 | 0.00017 |
|  | MMP9 | 17395 | 3.13 | 0.00055 |
|  | Defensin/DEF-A | 13240 | 3.13 | 0.00077 |
|  | IL-18rap/IL18-Rbeta | 16174 | 3.13 | 0.000040 |
|  | SLC11A/NRAMP | 18173 | 3.13 | 0.000040 |
|  | ITGB2/Pactolus | 16415 | 3.13 | 0.00017 |
|  | Nqp | 18054 | 3.13 | 0.00057 |
|  | S100A9/Calgranulin | 20202 | 2.64 | 0.000030 |
|  | PSGL1/p-selectin ligand | 20345 | 2.61 | 0.000050 |
| Immune Development | Pou2/Oct2 | 18987 | 2.32 | 0.000040 |
|  | CD300 | 246746 | 2.32 | 0.00011 |
|  | CD300a | 217303 | 2.04 | 0.000040 |
|  | FOS/AP1 | 14281 | 2.32 | 0.00025 |
|  | SYK-beta | 20963 | 2.32 | 0.000075 |
|  | TTC-7 | 225049 | 2.32 | 0.000075 |
|  | CD11B | 16409 | 2.32 | 0.00036 |
|  | LTB/P33 | 16994 | 2.32 | 0.000040 |
|  | dALA-S2/ASB | 11656 | 2.32 | 0.000075 |
|  | Sfpi-1 | 20375 | 2.32 | 0.000075 |
|  | VAV-1 | 22324 | 2.32 | 0.000040 |
|  | LIR3 | 18733 | 2.32 | 0.00025 |
|  | FYB | 23880 | 2.32 | 0.000036 |
|  | LIL-rb3 | 18733 | 2.32 | 0.000040 |
|  | NCF4/PHOX-P4 | 17972 | 2.32 | 0.00055 |
|  | PI3K-delta | 18707 | 2.32 | 0.00025 |
|  | PI3K-ap1 | 83490 | 2.32 | 0.00025 |
|  | PI3K-r5 | 320207 | 2.32 | 0.00025 |
| Immunoglobulin Related | TREM3/TLT3 | 58218 | 2.04 | 0.00038 |
|  | TREM4/TLT4 | 114840 | 2.04 | 0.00038 |
|  | FCG-R4 | 246256 | 2.04 | 0.00038 |
|  | TREML6 | 320148 | 2.04 | 0.00043 |
|  | PIL-Rbeta2 | 381678 | 2.04 | 0.00073 |

**TABLE S4.** Summary of microarray data for genes up-regulated 24 hrs after LPS insufflation.

| Group | Gene Name | I.D. | Enrichment Score | p-Value |
| --- | --- | --- | --- | --- |
| Oxidative Phosphorylation | ND1/NADH-Ubiquinone Dehydrogenase | 17716 | 29.93 | 1.3E-15 |
|  | ND2/NADH-Ubiquinone Dehydrogenase | 17717 | 29.93 | 1.3E-15 |
|  | ND3/NADH-Ubiquinone Dehydrogenase | 17718 | 29.93 | 1.3E-15 |
|  | ND4/NADH-Ubiquinone Dehydrogenase | 17719 | 29.93 | 1.3E-15 |
|  | ND6/NADH-Ubiquinone Dehydrogenase | 17722 | 29.693 | 1.3E-15 |
|  | CYTB/Cytochrome b | 17711 | 29.93 | 1.3E-15 |
|  | COX1 | 17708 | 29.93 | 1.3E-15 |
|  | COX2 | 17709 | 29.93 | 1.3E-15 |
|  | COX3 | 17705 | 29.93 | 1.3E-15 |
|  | COX 7A1 | 12865 | 29.93 | 1.3E-15 |
|  | 22 Trn Genes/tRNA Translation Elongation | 17726-17747 | 29.93 | 1.3E-40 |
|  | Rnr1/Ribosomal RNA precursor | 17724 | 29.93 | 1.3E-40 |
|  | Rnr2/Ribosomal RNA precursor | 17725 | 29.93 | 1.3E-40 |
|  | mtTFB2/TFB2M | 15278 | 29.93 | 1.3E-40 |
|  | TST/mitochondrial thiosulfate transferase | 22117 | 3.47 | 8.9E-10 |
| Immunoglobulin/Antigen Binding | Igh2/IgA | 16061 | 3.73 | 0.000038 |
|  | Igh3/IgG2b | 16016 | 3.73 | 0.000038 |
|  | Igh6/IgM | 16019 | 3.73 | 0.000038 |
|  | IghC1 | 16142 | 3.73 | 0.000038 |
| Cell Surface/Signaling | CD59a | 12509 | 1.76 | 0.0076 |
|  | CD209 | 170786 | 1.76 | 0.0029 |
|  | Hhip | 15245 | 1.76 | 0.0076 |
|  | SLC6A2/NAT1 | 20538 | 1.76 | 0.0076 |
|  | ENPP1/BDNP1 | 18605 | 1.76 | 0.0076 |
|  | CD150/SLAMF1 | 27218 | 1.76 | 0.0076 |
| Adhesion/Migration | CX3CR1/ FractalkineR1 | 13051 | 1.76 | 0.0029 |
|  | CD3g | 12502 | 1.76 | 0.0029 |
|  | Lyz1/Lysozyme | 17110 | 1.76 | 0.0029 |
| Inflammation | IFNalpha | 15965 | 1.76 | 0.0029 |
|  | HYAL/NAT6 | 15586 | 1.76 | 0.0029 |

**TABLE S5.**
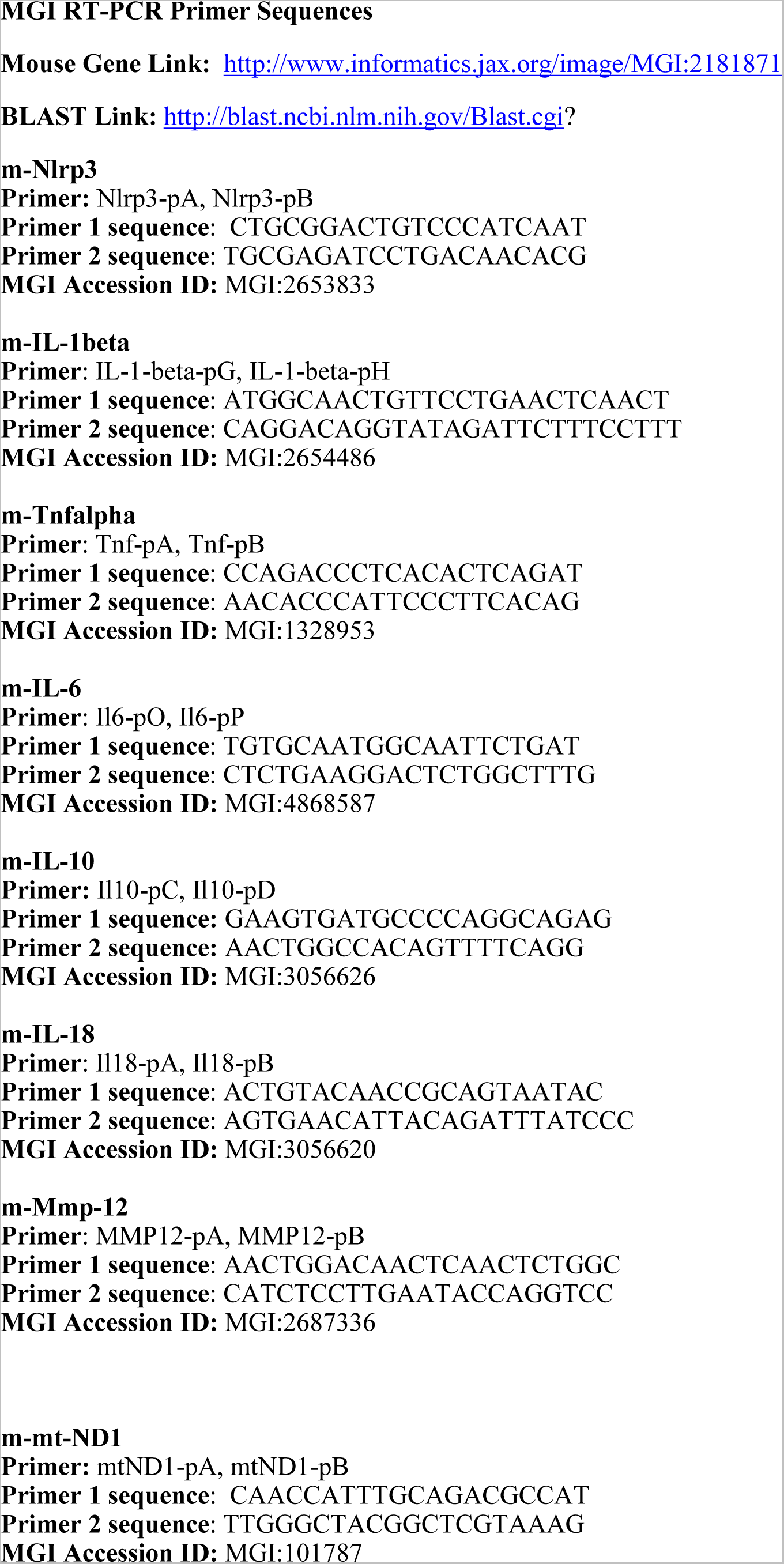
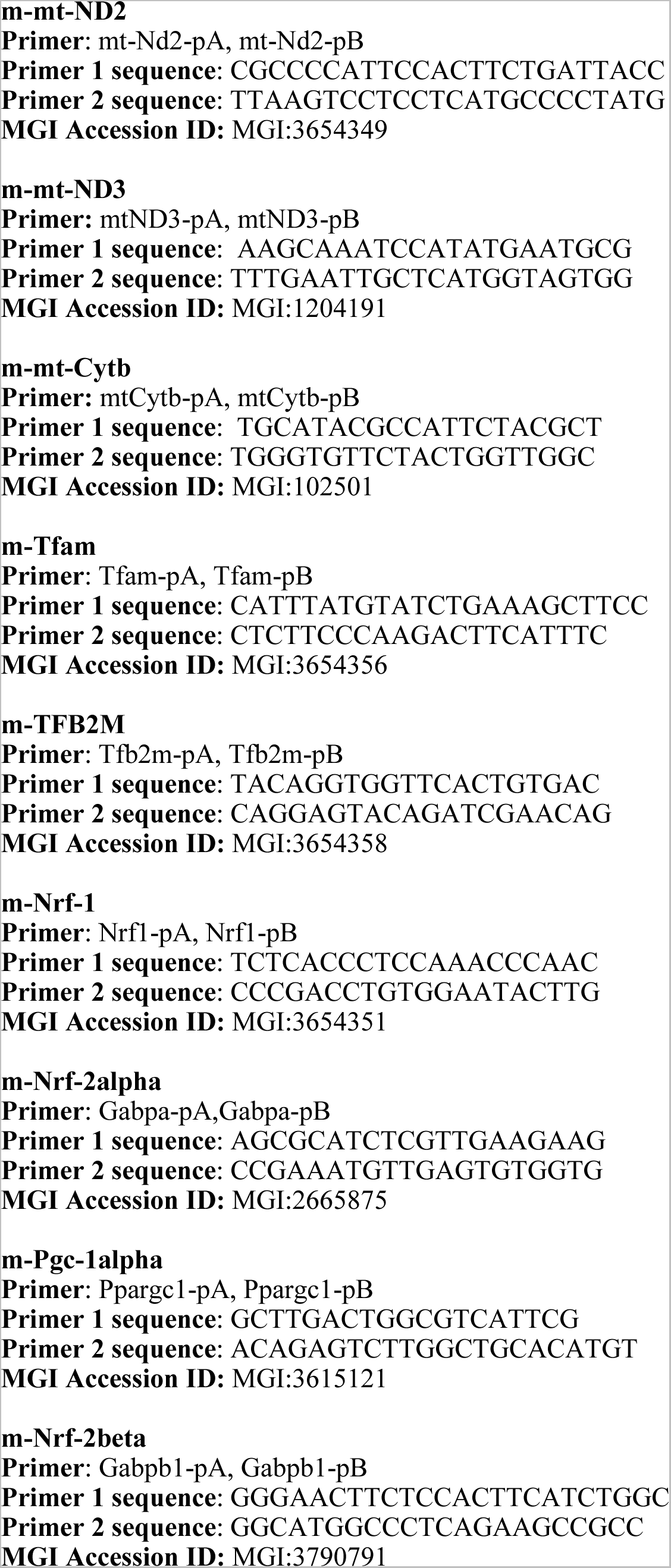
Primer sequences used for qRT-PCR analysis.

**TABLE S6.** Antibodies and sources.

| **Antibodies Used** |  |  |
| --- | --- | --- |
| Antibody | Source | Catalog Number |
| IκBα Phospho-S32/36 | Santa Cruz | ab12135 |
| COX-II | AbCam | ab110258 |
| XOR | AbCam | ab109235 |
| Nrf2β | AbCam | ab31163 |
| IL-18 (C18) | Santa Cruz | sc-6177 |
| Caspase-1 p10 | Santa Cruz | sc-514 |
| PGC-1α | Santa Cruz | sc-13067 |
| ASC (N-15) | Santa Cruz | sc-22514-R |
| Galectin-3 | Santa Cruz | sc-32790 |
| NLRP3 | Novus | NBP2-12446 |
| IL-1β | R&D Systems | AF-401-NA |
| TFB2M | Lifespan Bioscience | LS-C139799-100 |
| CD11b | e-Bioscience | 53-0112-82 |
| CD11c | e-Bioscience | 53-0114-82 |
| p65 | Santa Cruz | sc-109 |
| Keap-1 | AbCam | ab66620 |
| F4/80 | e-Bioscience | 53-4801-82 |
| p50 | Santa Cruz | sc-114 |
| MnSOD | EMD-Millipore | 06-984 |
| β-Actin | Sigma | A2066 |

